# Phylogenetic analysis reveals how selection and mutation shape the coevolution of mRNA and protein abundances

**DOI:** 10.1101/2024.07.08.602411

**Authors:** Alexander L. Cope, Joshua G. Schraiber, Matt Pennell

## Abstract

The regulatory mechanisms that shape mRNA and protein abundances are intensely studied. Much less is known about the evolutionary processes that shape the relationship between these two levels of gene expression. To disentangle the contributions of mutational and selective processes, we derive a novel phylogenetic model and fit it to multi-species data from mammalian skin tissue. We find that over macroevolutionary time: 1) there has been strong stabilizing selection on protein abundances; 2) mutations impacting mRNA abundances have minimal influence on protein abundances; 3) mRNA abundances are under selection to track protein abundances, and 4) mRNA abundances adapt more quickly than protein abundances due to increased mutational opportunity. We find additional support for these findings by comparing gene-specific parameter estimates from our model to human functional genomic data. More broadly, our new phylogenetic approach provides a foundation for testing hypotheses about the processes that led to divergence in gene expression.

## Introduction

Evolutionary divergence in gene expression is a major contributor to phenotypic divergence [1]. Accordingly, many studies leveraged high-throughput DNA microarrays and RNAseq to investigate how evolutionary processes shape patterns of mRNA abundances [2–6]. However, mRNA abundances provide an incomplete picture of gene expression evolution. Studies across a small but diverse set of taxa found differences in mRNA abundances need not imply differences in protein abundances [7–11]. Indeed, a fundamental question in molecular biology is understanding how various regulators and associated molecular mechanisms — from transcription to protein degradation — act in concert to produce the necessary amount of protein of a given gene for a cell to function [12, 13]. Despite our improved understanding of the molecular basis of this regulation, this does not provide a complete answer to the evolutionary question as to how and why these regulatory mechanisms themselves evolve.

To do that, we need other lines of evidence. First, there is the precise degree of correlation between steady-state abundances of mRNA and protein abundances across different genes, and how this correlation varies across organisms; while there is plenty of debate around the exact numbers, it is broadly true that the correlation is positive and strong, but imperfect [14, 15]. Second, genetic variants associated with changes in expression of mRNA (i.e., eQTLs) or proteins (i.e., pQTLs) are often distinct [16–18]. Second, comparative studies revealed that protein abundances are generally more conserved than mRNA abundances [7, 8, 19]. Third, there is some evidence for an evolutionary buffering effect in which evolutionary changes at one level of regulation are offset by subsequent changes to another level of regulation, ultimately leading to little change in protein abundances [9–11, 20]. This could be due to off-setting changes in transcription and translation, as has been documented in yeasts [9, 20], or between transcription and post-translational regulation (i.e., degradation), as has been found in primates [10, 16]. The mechanisms involved in such evolutionary buffering may, or may not, be distinct from the buffering mechanisms that may keep protein levels relatively stable despite transcriptional bursts and other causes of “noise” in gene expression [21–23].

An intuitive appealing explanation of this evidence is that of “compensatory evolution”, in which protein abundances, being “closer” to the phenotype than mRNA abundances, are generally under stabilizing selection; regulatory mutations that push proteins away from this optimum can fix if they are compensated by other types of regulatory mutations that move the protein abundances back. However, this model is inconsistent with the changes that occurred during a long-term experimental evolution study [24]. The model also implicitly predicts that mRNA abundances should show interspecific divergences that look like those produced by neutrality [25], which is also inconsistent with abundant comparative evidence that mRNA abundances are highly constrained [6, 26, 27]. And critically, being a purely verbal model, the compensatory evolution explanation neither makes any predictions for how much mRNA and protein abundances should diverge from one another (i.e., at what observed level of correlation would this explanation no longer be valid?) nor the relative importance of various evolutionary or regulatory processes in shaping patterns of gene expression across lineages.

To gain these richer insights, we need an explicit statistical model of the evolutionary and regulatory processes. To this end, we derived a set of novel phylogenetic models, rooted in first evolutionary and molecular principles, as well as Bayesian Markov Chain Monte Carlo machinery for fitting this model to comparative data. By comparing the fit of alternative models to matched transcriptome and proteome datasets from mammalian skin cell samples taken from ten species and by examining the values of estimated parameters, our model reveals new insights into the sources of evolutionary divergence in gene expression. For the sake of tractability, we had to make some strong, simplifying assumptions about the complex evolutionary processes that led to patterns of gene expression divergence. As such, there may be alternative interpretations of the parameter values we estimated (i.e., if we left out some key process, its effect may be included in the estimates of parameters we did include; [6]). However, our interpretations do make some clear predictions, which we tested using complementary functional genomic data from humans, mice, and fruit flies, which were not among the ten species we analyzed — these data thus provide critical out-of-sample tests of our model.

### A macroevolutionary model of mRNA and protein abundance coevolution

An overall schematic of our model is provided in Figure 1. Our model is founded on the basic premise that there are two possible types of mutations. First, there are “mRNA mutations” that influence the steady-state mRNA abundance and, in turn, impact the downstream protein steady-state abundances; this includes both mutations that influence transcription and those that impact mRNA degradation. These mutations occur at rate *µ*_*R*_ per genome per generation and are governed by parameters *c*, a constant that relates changes in mRNA to changes in protein, and 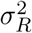, which quantifies the distribution of mutational effects on mRNA abundance. The parameter *c* can be interpreted as the degree to which mRNA mutations are buffered (|*c*| < 1) or amplified (|*c*| > 1) at the protein level.

**Figure 1:**
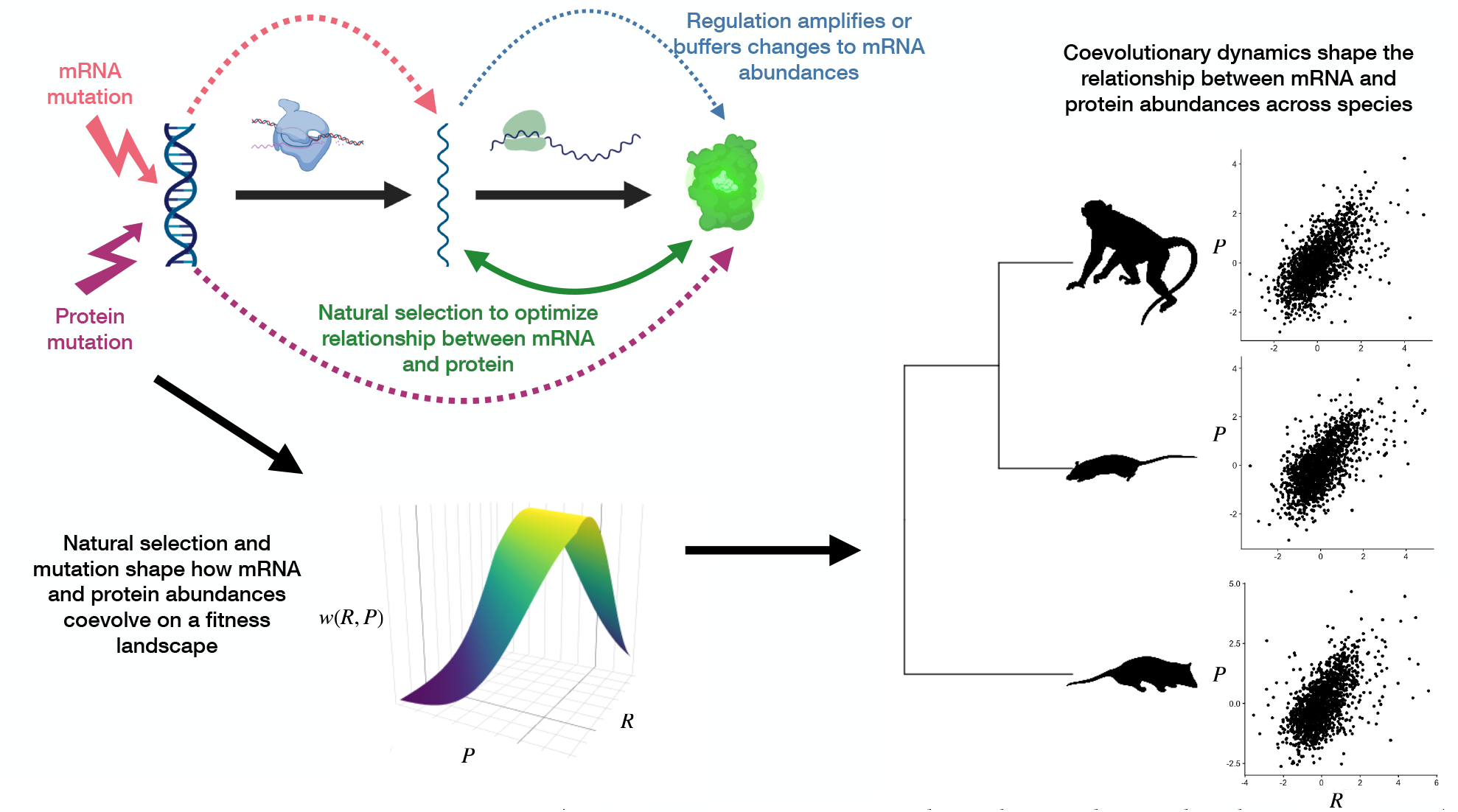
Conceptual framework of phylogenetic model. Scatter plots show relationship between mRNA and protein abundances are shown for *Macaca mulatta, Rattus norvegicus*, and *Monodelphis domestica* from [19]. Icons for DNA, RNA polymerase, mRNA, ribosome, and protein were obtained from BioRender.com. Images of species were taken from Phylopic (https://www.phylopic.org/).

Second, there are “protein mutations” that impact protein abundance (e.g., mutations that influence protein degradation); importantly, these do not directly influence steady-state mRNA abundances. These mutations occur at rate *µ*_*P*_ per genome per generation and depend on the parameter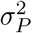, which quantifies the distribution of mutational effects on protein abundance that do not impact mRNA abundance.

To model the fitness of given mRNA and protein abundances, we wish to capture the idea that there may be a cost to having mRNA and protein abundances that are “mismatched” [28, 29]. To capture this, we model the optimal mRNA abundance as a linear function of protein abundance, 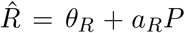 and the optimal protein abundance as a linear function of mRNA abundance, 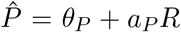. We use a Gaussian fitness function, as is standard in the evolutionary quantitative genetics literature [30], where the strength of selection *V*_*R*_ and *V*_*P*_ measures the width of the fitness function. Note that smaller *V*_*i*_ indicates stronger selection, as the fitness function becomes more peaked. (Here and elsewhere, we use the subscript *i* to indicate cases where we are referring to both the mRNA and protein analogs of a parameter.)

In the Supplementary Materials, we incorporate the above mutational and fitness models into a strong selection, weak mutation framework in which deleterious or beneficial mutations are either expunged from or fixed in diploid population of size *N* before another one arises [31]. This allows us to derive a pair of coupled stochastic differential equations describing the joint evolution of steady-state mRNA and protein abundances subject to mutation and selection. We find that compound parameters describing the combined effects of mutation, selection, and drift are important. First 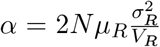 describes the rate of adaptation of the mRNA. Second, 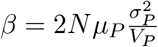 describes the rate of adaptation of the protein. Note that in both cases, the rate of adaptation is a function of mutational target size (as indicated by the *µ*_*i*_), mutational effect size (as indicated by the *σ*_*i*_), and the strength of selection (as indicated by the *V*_*i*_). Mutation and genetic drift also influence evolution through the parameters 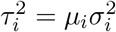.

The most general model, in which all the parameters can take arbitrary values, can generate complex feedback loops between *R* and *P*. Such a model is both computationally intractable and very difficult to interpret. We therefore focused on three limiting cases of our general model. These cases represent different ends of the spectrum of possible evolutionary scenarios, and we can compare them using model selection techniques. In all of these models, we describe the dynamics using a diffusion process, where B_*t*_ is uncorrelated two-dimensional Brownian motion.

In the first model, we assume that mRNA mutations play an important role on macroevolutionary timescales, and that natural selection acting on mRNA abundances is substantially stronger than natural selection acting on those of proteins. We refer to this model as “mRNA-driven evolution” and in the Supplementary Materials, we show that this model can be written

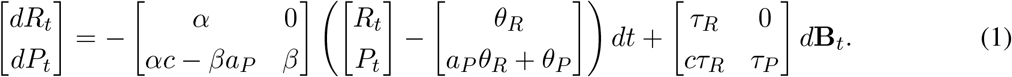

In the second model, mRNA mutations have a negligible impact on evolutionary changes to protein abundances, with evolutionary changes to mRNA abundances largely responding to changes to protein abundances. Here, we set *c* = 0, this would imply the effects of mRNA mutations are buffered, suggesting the existence of regulatory mechanisms downstream of mRNA abundances to leave protein abundances largely unaltered. We refer to this model as “protein-driven evolution” and show that this can be written as:

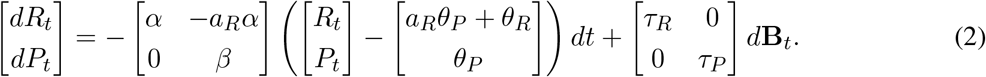

Under both the mRNA-driven and protein-driven models, we find that mRNA and protein levels are correlated (see Supplementary materials). Under the mRNA-driven model, the strength of correlation is determined linearly by *c* and *a*_*P*_, while under the protein-driven model, it is determined linearly by *a*_*R*_.

Last, as a null model, we considered that the evolutionary dynamics of mRNA and protein abundance are uncoupled. We refer to this model as “independent evolution”. Then, the model can be written

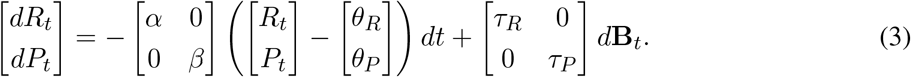

These are all coupled Ornstein-Uhlenbeck (OU) processes, whose likelihood can be computed efficiently on a phylogeny [32]. Univariate OU processes (in which, like our null model, adaptation in one variable does not depend on any other measured variable) have been widely applied to describe the evolution of mRNA counts along a phylogeny [6, 26] but to our knowledge, this is the first application of a coupled OU processes to study gene expression evolution at two different levels simultaneously. In order to fit special cases of our model using a Bayesian approach, we developed a custom Markov Chain Monte Carlo (MCMC) machinery for estimating the parameters. Using extensive simulations (see Supplementary Materials), we validated that our estimation procedure was able to recover the generating parameters under a range of conditions and that standard model selection techniques [e.g., 33] are able to correctly identify the generating scenario.

### Evolution of gene expression in mammalian fibroblasts skin tissue is determined by selection on protein abundance

We analyzed a recently published dataset [19] of mRNA and protein abundances measured for ten species of mammalian skin samples. Critically, these were all measured by the same group of researchers, using standardized experimental protocols to minimize the impact of technical variation. The phylogenetic tree of these ten species and the correlations between mRNA and protein levels are shown in Figure 2A; the pairwise similarity as a function of divergence time is shown in Figure 2B.

**Figure 2:**
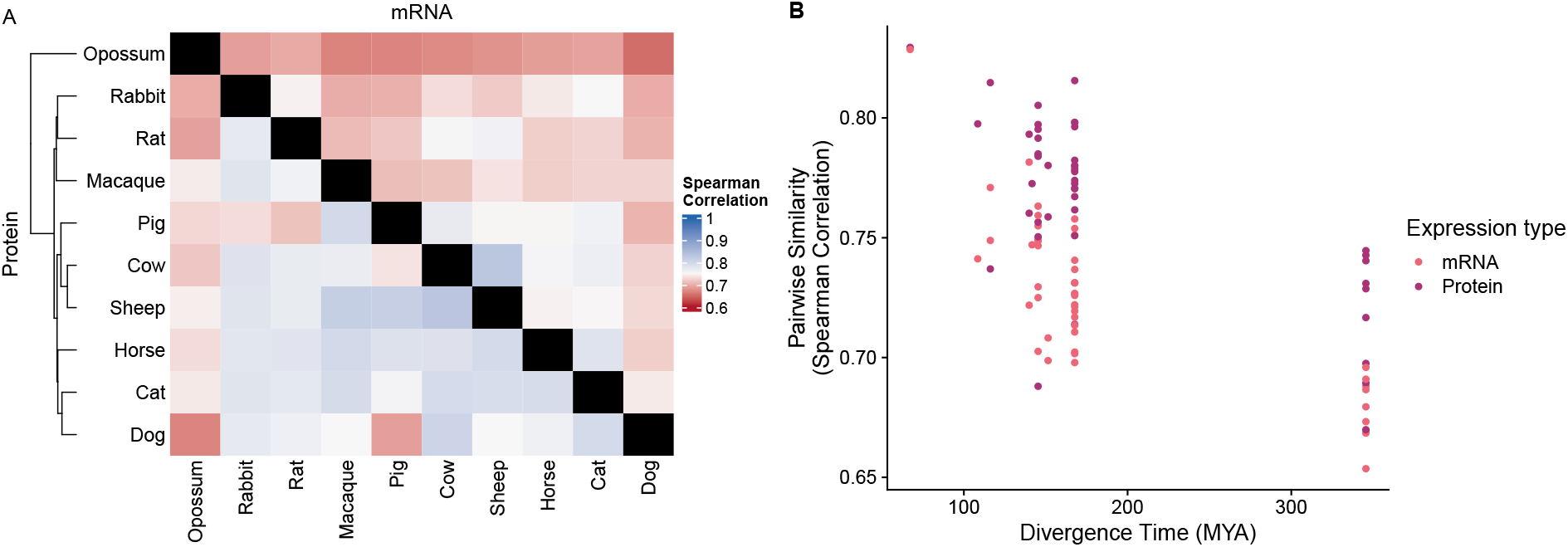
(A) Heatmap showing across-species pairwise comparisons (measured as the Spearman rank correlation) of mRNA (upper triangle) and protein abundances (lower triangle). The phylogenetic tree of the 10 species is included on the lower triangle. (B) Pairwise correlations of mRNA and protein abundance as a function of species divergence time.

Ten species is a large sample size for paired transcriptomic and proteomic data but nonetheless small relative to the number of parameters we sought to estimate. To overcome this limitation, we assumed that the steady-state mRNA and protein abundances for each of the 1,641 orthologous genes in the dataset were an independent outcome of the same co-evolutionary process. As such, we estimated unique steady-state optima for mRNA and protein abundances of each gene, both functions of the gene-specific parameters *θ*_*R*_ and *θ*_*P*_ ; all other parameters were assumed to be equal across genes. Although there is certainly variation in the co-evolutionary dynamics across genes, we demonstrated using simulations that our model parameter estimates closely match the mean values across genes (see Supplementary Materials).

Overall, we find overwhelming support for the protein-driven evolution model (Figure 3A): the DIC value for the protein-driven model is 4,201 DIC units lower than the next-best model. From this model comparison, we infer that mRNA mutations have a negligible impact on protein abundances over long evolutionary timescales (i.e., *c* is close to 0; even under the disfavored mRNA-driven model, we estimate *c* to be 0.141 (95% CI: 0.079 – 0.197)). There are many mechanisms by which cells minimize the impact of variation in mRNA abundances on protein production [12, 13]; our data suggest these mechanisms influence the macroevolutionary dynamics of steady-state expression and not just the cellular states.

**Figure 3:**
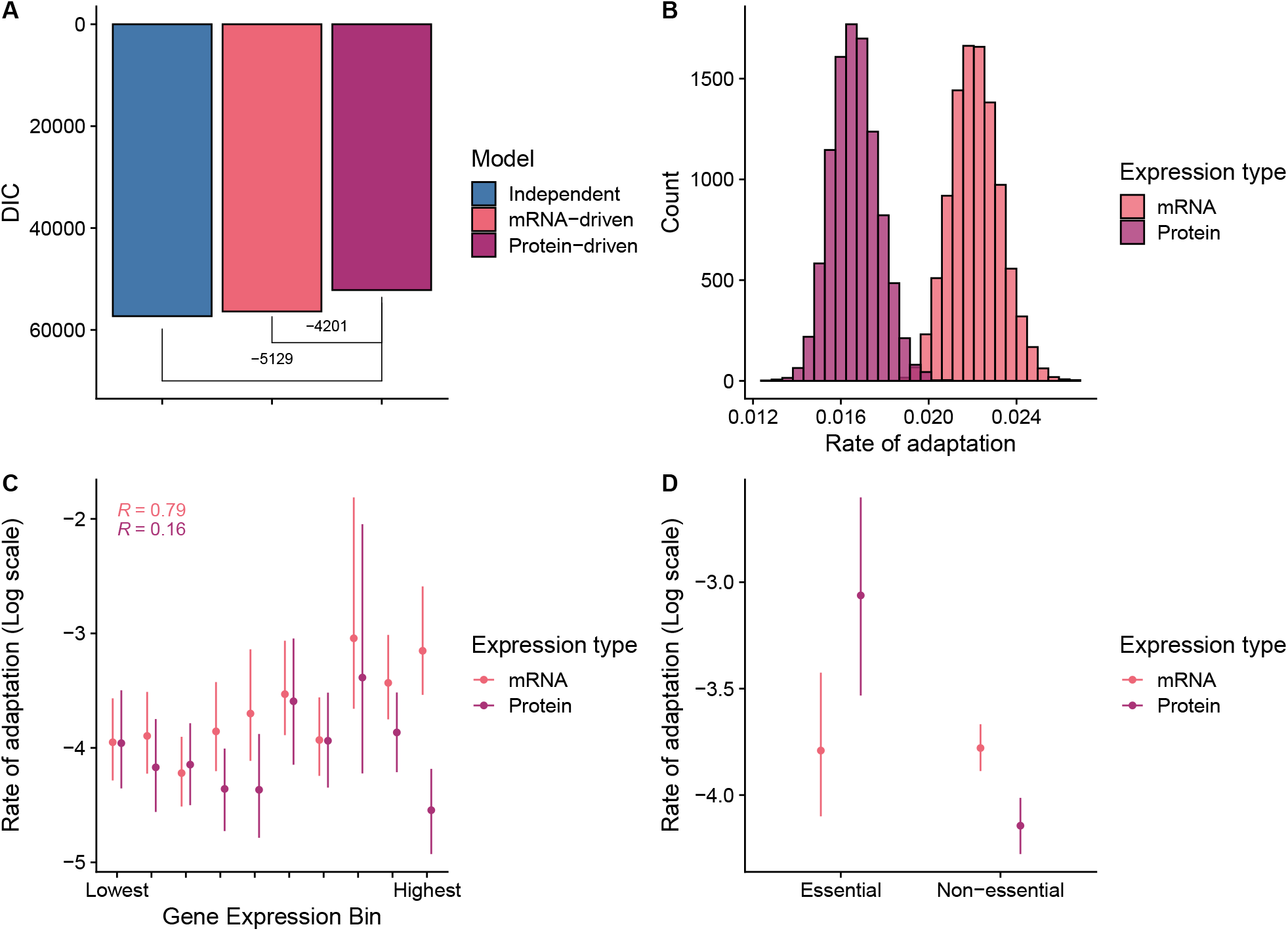
Comparisons of models and posterior distributions of model parameters from protein-driven model fit. (A) Comparison of DIC values for protein-driven, mRNA-driven and, independent models. The protein-driven model has the lowest DIC value, indicating it is the best of the 3 models. Brackets indicate the ΔDIC values relative to the protein-driven model. (B) mRNA and protein abundance rate of adaptation parameters *α* and *β*, respectively. (C) Rate of adaptation parameter *α* (mRNA) and *β* (protein) across gene expression decile bins. Deciles were determined using the integrated protein abundance data for humans from PaxDB. (D) Rate of adaptation parameters across “Essential” and “Non-essential” genes in humans *in vitro* cell lines as previously defined [38].

### mRNA abundances adapt more rapidly than protein abundances

Examining the parameter values estimated from the preferred protein-driven evolution model, we find that the two parameters describing the rate of adaptation, *α* and *β*, have posterior means of 0.022 (95% Credible Interval (CI): 0.019–0.024) and 0.016 (95% CI: 0.014–0.018), respectively (Figure 3B). These estimates correspond to a mean “phylogenetic half-life” [34] of 31 Million Years (MY) (95% CI: 28–34 MY) for the mRNA abundances and 41 MY (95% CI: 36–46 MY) for the protein abundances. This means that it takes longer for the proteins to adapt to changes in their optima than it does for mRNA and is consistent with previous observations that protein abundances are more similar between species than mRNA abundances [7, 8, 19].

multiple lines of evidence indicate natural selection is stronger on highly expressed genes [35, 36]. If our model is actually capturing the proposed evolutionary processes, we predict adaptation parameters *α* and *β* are correlated with expression level. To test this prediction, we binned genes into deciles based on an integrated measure of protein abundances in humans taken from PaxDB 5.0 [37]. We then re-fit the preferred protein-driven model to the binned data. We found that genes that were more highly expressed tended to have stronger rate of adaptation parameters for both mRNA and protein abundances, except for the highest expression categories in the case of proteins, in which the direction of the trend reversed (Figure 3C). (We do not calculate *P*-values as the sample size is a property of the binning strategy; we divided the data into gene expression deciles to ensure that we could still estimate parameters within each set of binned genes.)

We further validated our inferences by testing whether genes whose functional roles make them likely to be under strong purifying selection indeed have stronger selection terms estimated from our model (Figure 3D). Using annotations from [38], we found that, as predicted by our protein-driven model, the rate of adaptation for protein abundances was significantly stronger in essential genes (on natural scale, *β* = 0.048, 95% CI: 0.027 – 0.071) versus non-essential genes (on natural scale, *β* = 0.015, 95% CI: 0.013 – 0.017). There was no difference between the selection on mRNA between the essential (on natural scale, *α* = 0.0229, 95% CI: 0.015 – 0.030) and non-essential genes (on natural scale, *α* = 0.0228, 95% CI: 0.020 – 0.025), consistent with our inference that natural selection on protein levels is the primary driver of gene expression evolution.

### Genetic architecture is predictive of macroevolutionary trends

Based on mathematical reasoning described in depth in the Supplementary Material, our model predicts that the rate of adaptation is influenced by mutational input. Although it is difficult to quantify mutational input to mRNA and protein abundances, we used several proxies to assess the predictions of our model. First, we hypothesized that the number of *cis*-regulatory elements is correlated with mutational target size for mRNA abudnances, but not protein abundances. We obtained enhancer information from the EnhancerAtlas2.0 database [39] to estimate the number of active enhancers per gene in human keratinocyte cells, the primary cell type found in the epidermis. To control for confounding due to the absolute value of gene expression, we calculated the mean number of enhancers per gene for each expression bin and compared this average to the estimated parameter value of each bin. Consistent with our interpretation, genes with more enhancers in human keratinocyte cells show a faster rate of adaptation at the mRNA levels (Spearman rank correlation *R* = 0.94, Figure 4B) but not the protein level. We recapitulated this pattern using enhancer counts from mouse cells, but could not replicate it using *Drosophila melanogaster* enhancer counts (see Supplementary Materials), showing that our result likely reflects true dynamics within mammals that are not conserved at long evolutionary distances.

**Figure 4:**
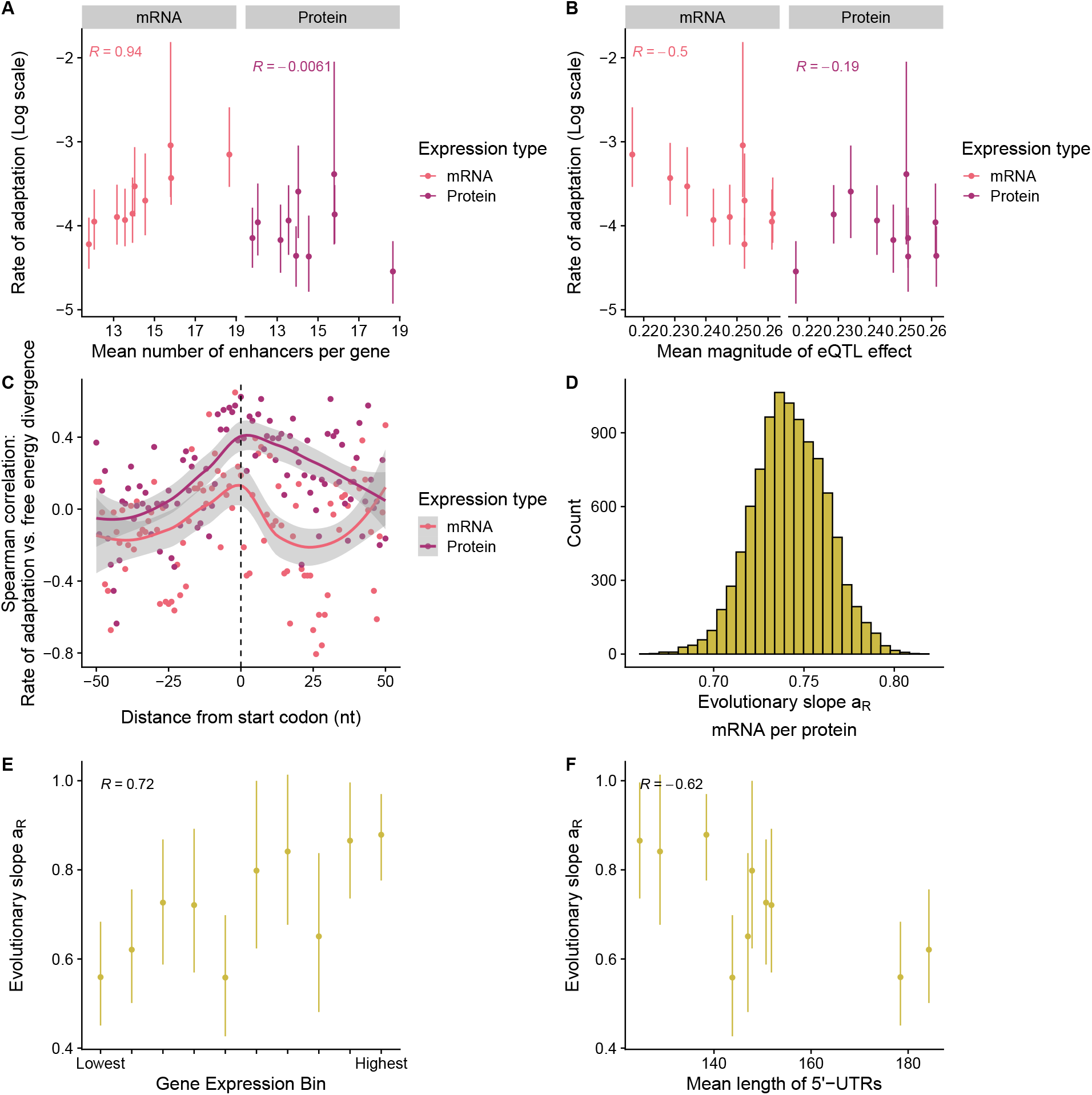
Validation of model parameter estimates using out-of-sample human functional genomics data. Fits were based on protein-driven model. All error bars represent 95% credible intervals. Spearman rank correlations *R* are reported. (A) The relationship between the mean number of enhancers per gene and the rate of adaptation for mRNA and protein abundances. Rate of adaptation estimates are taken from the decile bins based on gene expression. (B) Similar to (A), but using the mean absolute magnitude of eQTL effects in humans obtained from GTEx. (C) The relationship between divergence in mRNA secondary structure (specifically, free energy) near the start codon for human and mouse genes [42] and the rates of adaptation. (D) Posterior distribution of the evolutionary slope parameter *a*_*R*_. (E) Variation in *a*_*R*_ across gene expression decile bins. (F) Relationship between the mean length of human 5’-UTRs and *a*_*R*_.

As an alternative measure of mutational input for mRNA abundance, we examined eQTLs in humans using data from GTEx (version 8) [40]. We calculated the mean absolute effect size of eQTLs for each gene across all available human datasets in GTEx, which includes multiple tissue types. As with enhancers, we then calculated the mean absolute eQTL effect size across all genes within each gene expression decile, as before. Surprisingly, we find that genes with larger eQTL evolve under weaker selection (Spearman rank correlation *R* = −0.5, Figure 4D). We speculate this is because eQTLs need to be both common enough and have large enough effects to be detected using genome-wide association study methods. Thus, eQTL for genes under strong selection must have small effects, while large effect eQTL must impact genes under weak selection. Our observation here aligns with the recent discovery that there is little overlap between genes with eQTL and genes associated with disease due to this trade-off between effect size and strength of selection [41]. We see a similar, albeit buffered, impact on protein adaptation, consistent with our inference that mRNA mutations are buffered over macroevolutionary timescales (Spearman rank correlation *R* = −0.19).

On the other hand, cells can regulate protein abundance independently of mRNA abundance by tuning mRNA translation, particularly the ability of ribosomes to initiate translation. Initiation efficiency is regulated by numerous transcript features, including mRNA secondary structure near the CDS start codon and properties of the 5’-untranslated region (5’-UTR). Using previously estimated mRNA secondary structure ensemble free energies near the CDS start codon for human and mouse [42], we found that divergence in secondary structure between human and mouse is associated with higher rates of adaptation for protein, but not mRNA, abundance (Figure 4C). The correlation between the rate of adaptation and divergence of mRNA secondary structure peaks for protein abundances near the start codon, consistent with changes to translation initiation efficiency significantly altering protein abundances [43]. Consistent with the importance of the 5’-UTR as a *cis* regulator of translation, we found that human genes with less conserved 5’-UTRs (based on PhyloP scores taken from [44]) showed higher rates of adaptation in protein abundances (see Supplementary Materials, Spearman rank correlation *R* = −0.45). In contrast, genes with higher rates of adaptation for mRNA abundances generally showed greater conservation of 5’-UTRs (Spearman rank correlation *R* = 0.15). These results indicate that the evolution of regulators of mRNA translation plays a significant role in shaping the evolution of steady-state protein abundances independent of mRNA abundances. While changes to mRNA translation can alter mRNA degradation rates [45, 46], our results suggest changes to mRNA translation have a negligible effect on the evolution of steady-state mRNA abundances compared to protein abundances.

### Strong selection to maintain correlations between mRNA and protein abundances

The *a*_*R*_ parameter in the protein-driven model represents the linear strength of selection for the mRNA abundances to match the protein abundances; we estimated this parameter to be 0.742 (95% of CI: 0.702– 0.782, Figure 4D). The observation that *a*_*R*_ is less than but, close to, 1 suggests there is strong selection for the mRNA levels to match the protein levels but that there is a substantial evolutionary lag for the alignment to equilibrate following selection for changes to the protein levels. This highlights the value of building the relevant mechanisms into the statistical model itself because it is not obvious from looking at any particular level of correlation which mechanisms need to be invoked.

More highly-expressed genes have evolutionary slopes *a*_*R*_ closer to 1 compared to moderate and low expression genes (Spearman rank correlation *R* = 0.72, Figure 4E). This is consistent with the observation that highly-expressed genes tend to be more translationally efficient compared to moderate and low-expressed genes. To test if our model parameters generally captured the expected translation efficiency, we compared to our estimate of the optimal steady-state protein and mRNA abundances to TE estimated from independent human ribosome profiling and RNA-seq measurements [47]. We find that the optimum and human mean TEs per bin are highly-correlated (Spearman rank correlation *R* = 0.96, see Supplementary Materials). We next hypothesized that genes with more opportunity for mutations to impact protein abundance may not require mRNA and protein abundances to be as highly correlated. Confirming this hypothesis, we found that estimates of *a*_*R*_ are anti-correlated with the length of 5’-UTRs in humans (Spearman rank correlation *R* = −0.62, Figure 4F) and in mouse (Spearman rank correlation *R* = −0.64, see Supplementary Materials). Given that increasing the 5’-UTR length increases mutational input and thus the potential for changes to translational regulation, it is unsurprising that *a*_*R*_ is close to 1 for genes with shorter 5’-UTRs across species, indicating more importance for mRNA abundance to match protein abundance when there is less opportunity to regulate protein abundance independently of mRNA abundance.

## Conclusion

Numerous studies have used phylogenetic models to investigate the evolutionary processes that shaped variation in mRNA abundances among species [6]. In this paper, we take this work a big step forward and use phylogenetic models to infer the processes that have shaped the co-evolution between mRNA and protein abundances. Doing so required us to derive novel, mechanistic models from first principles to capture the biochemical and evolutionary relationships between these variables. We suggest that this general modeling framework can serve as the foundation for further investigations into the role of complex processes in shaping patterns of gene expression. Remarkably, we show the evolutionary parameters estimated from these models are consistent with within-species functional genomic information. In evolutionary biology more broadly, it has typically been difficult to map the parameters estimated from macroevolutionary to microevolutionary processes [48] — our findings imply that, at least for gene regulation, we may be able to understand the general rules that shape evolutionary dynamics across timescales.

## Data availability

No new data were generated for this study. The publicly-available data used in this study are available via the references provided in this manuscript. R notebooks and R scripts for recreating our analyses can be found at https://github.com/phylo-lab-usc/rna-protein-coevolution.

## Acknowledgements

We thank Rex Jiang, Qifan Wang, Melissa Guzman, and Rachel Brem for comments on the manuscript, as well as the Pennell, Edge, and Mooney lab groups for helpful discussion of this work. Fabio Machado and Josef Uyeda provided advice on our implementation of the MCMC. Tony Zheng, Jeffrey Spence, Hakhamanesh Mostafavi, and Jonathan Pritchard provided curated data on gene function. Gunter Wagner provided the phylogenetic tree associated with the expression data analyzed here. This work was supported by the NIH-funded Rutgers INSPIRE IRACDA Postdoctoral Program (grant #GM093854 to ALC) and NIH grant R35GM151348 to MP.

