## supplementary material for "Phylogenetic analysis reveals how selection and mutation shape the coevolution of mRNA and protein abundances"

4.2.3 Determining mutational input using the number of regulatory elements and eQTLs . 17
4.2.4 Quantifying gene properties related to mRNA translation and translation efficiency . 17

### 1 Derivation of phylogenetic model

#### 1.1 Mutational model

We model the (log) mRNA expression, $R$, and the (log) protein expression $P$. Broadly, we consider two types of mutations. First, there are mutations that impact mRNA abundances, and, via the central dogma, cascade to protein abundances. These mutations occur with probability $\mu_R$ per genome per generation and result in the outcome

$$R \to R' = R + \delta R$$
$$P \to P' = P + c\delta R$$

where $c$ is a constant that relates changes in mRNA to changes in protein, and $\delta R$ is a random variable with mean 0 and variance $\sigma_R^2$. On the other hand, if a mutation impacts protein abundances alone, which happens at rate $\mu_P$ per genome per generation, we have

$$R \to R' = R$$
$$P \to P' = P + \delta P.$$

where $\delta P$ is a random variable with mean 0 and variance $\sigma_P^2$

subsectionFitness model

We also specify a Gaussian fitnes function in which the optimal mRNA level is a linear function of the protein level, and similarly the optimal protein level is a linear function of the optimal mRNA level,

$$w(R,P) \propto \exp\left(-\frac{(P-\theta_P-a_P R)^2}{2V_P}-\frac{(R-\theta_R-a_R P)^2}{2V_R}\right).$$

Note that if we set $a_p \to 0$, then the protein has an optimal level that is independent of mRNA expression, and similarly if we set $a_R \to 0$, the mRNA has an optimal level that is independent of protein expression.

#### 1.2 Evolutionary dynamics

To derive the evolutionary dynamics under this model, we can use a strong selection, weak mutation (SSWM) limit [1]. In this framework, we assume that there is only one segregating mutation that impacts either protein or mRNA abundances at any given time.

Assuming that $\sigma_i^2$ are both small (i.e. mutations have small effects), we compute the selection coefficient of a mutation as

$$s \approx \log\left(\frac{w(R', P')}{w(R, P)}\right).$$

Write the impact of a mutation on mRNA level as $M_R$ and the impact on protein level as $M_P$ (so that for an mRNA mutation, $M_R = \delta R$ and $M_P = c\delta R$ while for a protein mutation, $M_R = 0$ and $M_P = \delta P$). Then, we approximate that both $M_R$ and $M_P$ are small so we discard terms with $M_R^2$, $M_P^2$ and $M_R M_P$ so that we can write

$$\begin{aligned} s &\approx M_R\left(a_P\frac{P - a_P R - \theta_P}{V_P} - \frac{R - a_R P - \theta_R}{V_R}\right) + M_P\left(a_R\frac{R - a_R P - \theta_R}{V_R} - \frac{P - a_P R - \theta_P}{V_p}\right) \\ &= M_R\frac{\partial}{\partial R}\log(w(R,P)) + M_P\frac{\partial}{\partial P}\log(w(R,P)) \end{aligned}$$

where the first term is the fitness consequence from the impact the mutation has on mRNA abundances and the second term is the impact it has on protein abundances.

We then need to calculate the mean and variance in trait change following a fixation, denoted $\Delta X_i$, where $X_1 = R$, and $X_2 = P$. Given a mutation with selection coefficient $s$, we have that

$$\begin{aligned} \pi(s) &= \frac{1 - e^{-2s}}{1 - e^{-4Ns}} \\ &\approx \frac{1}{2N} + \left(1 - \frac{1}{2N}\right)s, \end{aligned}$$

where the approximation follows from assuming $s$ is small.

Then, following fixation we can use Bayes rule to write

$$p(M_R, M_P \mid \text{fix}) = \frac{\pi(s(M_R, M_P))p(M_R, M_P)}{\int\int \pi(s(M_R, M_P))p(M_R, M_P)d(M_R)d(M_P)}.$$

Note however that $\pi$ is a linear function of $s$, and $s$ is a linear function of the $M_i$. So the integral on the bottom is simply

$$\begin{aligned} &\int\int \pi(s(M_R, M_P))p(M_R, M_P)d(M_R)d(M_P) \\ &= \int\int\left(\frac{1}{2N} + \left(1 - \frac{1}{2N}\right)\left(M_R\frac{\partial}{\partial R}\log(w(R,P)) + M_P\frac{\partial}{\partial P}\log(w(R,P))\right)\right)p(M_R, M_P)d(M_R)d(M_P) \\ &= \frac{1}{2N} + \left(1 - \frac{1}{2N}\right)\left(\mathbb{E}(M_R)\frac{\partial}{\partial R}\log(w(R,P)) + \mathbb{E}(M_P)\frac{\partial}{\partial P}\log(w(R,P))\right) \\ &= \frac{1}{2N}, \end{aligned}$$

where the second line follows from linearity of the integral operator and recognizing an expectation, and the third line follows because we assume those expectations are 0. Intuitively because we model effect sizes as small and symmetrical, a typical mutation looks approximately neutral.

Thus, the distribution of mutations that fix is approximately

$$p(M_R, M_P \mid \text{fix}) \approx 2N\left(\frac{1}{2N} + \left(1 - \frac{1}{2N}\right)\left(M_R \frac{\partial}{\partial R}\log(w(R,P)) + M_P \frac{\partial}{\partial P}\log(w(R,P))\right)\right) p(M_R, M_P).$$

This distribution seems reasonable: it takes the underlying distribution of mutation effects and weights mutations that move up the fitness surface more highly.

To calculate the expected change, we compute

$$\begin{aligned}
\mathbb{E}(M_i \mid \text{fix}) &= \int M_i p(M_R, M_P \mid \text{fix}) dM_R dM_P \\
&= 2N\left(\frac{\mathbb{E}(M_i)}{2N} + \left(1 - \frac{1}{2N}\right)\left(\mathbb{E}(M_i M_R)\frac{\partial}{\partial R}\log(w(R,P)) + \mathbb{E}(M_I M_P)\frac{\partial}{\partial P}\log(w(R,P))\right)\right) \\
&= (2N-1)\left(\mathbb{E}(M_i M_R)\frac{\partial}{\partial R}\log(w(R,P)) + \mathbb{E}(M_i M_P)\frac{\partial}{\partial P}\log(w(R,P))\right)
\end{aligned}$$

Similarly, to compute the (co)variance in change, we compute

$$\text{Cov}(M_i, M_j \mid \text{fix}) = \mathbb{E}(M_i M_j \mid \text{fix}) - \mathbb{E}(M_i \mid \text{fix})\mathbb{E}(M_j \mid \text{fix}).$$

However, we're working in a regime where $E(M_i)$ is small, so $E(M_i \mid \text{fix})E(M_j \mid \text{fix})$ will be very small, and we can discard that term. Then we only need to compute the mixed moment,

$$\begin{aligned}
\mathbb{E}(M_i M_j \mid \text{fix}) &= \int M_i M_j p(M_R, M_P \mid \text{fix}) dM_R dM_P \\
&= 2N\left(\frac{\mathbb{E}(M_i M_j)}{2N}\right. \\
&\quad \left. + \left(1 - \frac{1}{2N}\right)\left(\mathbb{E}(M_i M_j M_R)\frac{\partial}{\partial R}\log(w(R,P)) + \mathbb{E}(M_i M_j M_P)\frac{\partial}{\partial P}\log(w(R,P))\right)\right).
\end{aligned}$$

However, if we make the assumption that the distribution is marginally symmetric (i.e. $\mathbb{E}(M_i^3) = 0$), this will eliminate all the terms with the expectation of 3 variables. Hence, we are left with

$$\text{Cov}(M_i, M_j) = \mathbb{E}(M_i M_j)$$

In our model, the distribution of mutation effect sizes (conditional on a mutation happening at all) can be written

$$p(M_R, M_P) = \frac{\mu_R}{\mu_R + \mu_P} p(M_R)\delta(M_P - cM_R) + \frac{\mu_P}{\mu_R + \mu_P} p(M_P)\delta(M_R).$$

To see this, note that a mutation is an mRNA mutation with probability $\frac{\mu_R}{\mu_R + \mu_P}$, in which case the effect size has some distribution $p(M_R)$ and the protein level has to change by exactly $cM_R$, which is enforced by the dirac delta function. On the other hand, a protein mutation occurs with probability $\frac{\mu_P}{\mu_R + \mu_P}$, and the effect size is drawn from $p(M_P)$, while the mRNA change is forced to be 0 by the dirac delta function.

The change to mRNA abundances will be the posterior expectation, $\mathbb{E}(M_R \mid \text{fix})$. To compute this, we will need the *prior* expectations

$$\mathbb{E}(M_R^2) = \frac{\mu_R}{\mu_R + \mu_P}\sigma_R^2,$$

and

$$\mathbb{E}(M_R M_P) = c\frac{\mu_R}{\mu_R + \mu_P}\sigma_R^2$$

and

$$\mathbb{E}(M_P^2) = c^2\frac{\mu_R}{\mu_R + \mu_P}\sigma_R^2 + \frac{\mu_P}{\mu_R + \mu_P}\sigma_P^2,$$

which are all computed by directly integrating against the prior density.

Then, the expected change to mRNA abundances is

$$\begin{aligned}
\mathbb{E}(M_R \mid \text{fix}) &= (2N-1)\left(\mathbb{E}(M_R^2)\frac{\partial}{\partial R}\log(w(R,P)) + \mathbb{E}(M_R M_P)\frac{\partial}{\partial P}\log(w(R,P))\right) \\
&= (2N-1)\left(\frac{\mu_R}{\mu_R+\mu_P}\sigma_R^2\frac{\partial}{\partial R}\log(w(R,P)) + c\frac{\mu_R}{\mu_R+\mu_P}\sigma_R^2\frac{\partial}{\partial P}\log(w(R,P))\right) \\
&= (2N-1)\frac{\mu_R}{\mu_R+\mu_P}\sigma_R^2\left(\frac{\partial}{\partial R}\log(w(R,P)) + c\frac{\partial}{\partial P}\log(w(R,P))\right).
\end{aligned}$$

This seems reasonable, because for mRNA to change, you need an mRNA mutation, and it needs to weight both the effect on mRNA fitness AND the effect on protein fitness. If we fully evaluate the fitness functions, this results in

$$-(2N-1)\frac{\mu_R}{\mu_R+\mu_P}\sigma_R^2\left((1-a_Rc)\frac{R-a_RP-\theta_R}{V_R} + (c-a_p)\frac{P-a_PR-\theta_P}{V_P}\right).$$

Next, the expected change to protein abundances is

$$\begin{aligned}
\mathbb{E}(M_P \mid \text{fix}) &= (2N-1)\left(\mathbb{E}(M_R M_P)\frac{\partial}{\partial R}\log(w(R,P)) + \mathbb{E}(M_P^2)\frac{\partial}{\partial P}\log(w(R,P))\right) \\
&= (2N-1)\left(c\frac{\mu_R}{\mu_R+\mu_P}\sigma_R^2\frac{\partial}{\partial R}\log(w(R,P)) + \left(c^2\frac{\mu_R}{\mu_R+\mu_P}\sigma_R^2 + \frac{\mu_P}{\mu_R+\mu_P}\sigma_P^2\right)\frac{\partial}{\partial P}\log(w(R,P))\right) \\
&= (2N-1)c\frac{\mu_R}{\mu_R+\mu_P}\sigma_R^2\left(\frac{\partial}{\partial R}\log(w(R,P)) + c\frac{\partial}{\partial P}\log(w(R,P))\right) \\
&\quad + (2N-1)\frac{\mu_P}{\mu_R+\mu_P}\sigma_P^2\frac{\partial}{\partial P}\log(w(R,P)).
\end{aligned}$$

This makes sense: the first term is the expected change in mRNA, but reduced by $c$, since under our muational model, every mRNA mutation induces an instant change in protein level. Then, the second term accounts for changes that only impact protein level. Again, evaluating the fitness functions will result in

$$\begin{aligned}
&- c(2N-1)\frac{\mu_R}{\mu_R+\mu_P}\sigma_R^2\left((1-a_Rc)\frac{R-a_RP-\theta_R}{V_R} + (c-a_p)\frac{P-a_PR-\theta_P}{V_P}\right) \\
&- (2N-1)\frac{\mu_P}{\mu_R+\mu_P}\sigma_P^2\left(\frac{P-a_PR-\theta_P}{V_P} - a_R\frac{R-a_RP-\theta_R}{V_R}\right).
\end{aligned}$$

We also have immediately that

$$\mathrm{Var}(M_R \mid \text{fix}) = \mathbb{E}(M_R^2) = \frac{\mu_R}{\mu_R + \mu_P}\sigma_R^2,$$

and

$$\mathrm{Var}(M_P \mid \text{fix}) = \mathbb{E}(M_P^2) = c\frac{\mu_R}{\mu_R + \mu_P}\sigma_R^2 + \frac{\mu_P}{\mu_R + \mu_P}\sigma_P^2,$$

and finally

$$\mathrm{Cov}(M_R, M_P \mid \text{fix}) = \mathbb{E}(M_R M_P) = c\frac{\mu_R}{\mu_R + \mu_P}\sigma_R^2.$$

#### 1.3 Converting to an mvOU framework

The model described above is evidently a multivariate Ornstein-Uhlenbeck model, but to use it for inference, it's helpful to turn it into the standard form [2]

$$d\mathbf{X}_t = -\mathbf{F}(\mathbf{X}_t - \hat{\mathbf{X}})dt + \mathbf{\Sigma} d\mathbf{B}_t$$

Here, we use the convention that $R$ is indicated by the first entry in the $\mathbf{X}$ vector and $P$ is indicated by the second entry.

First, we need to move from the jump chain, outlined above which is happening at discrete steps when a mutation occurs, into a continuous time model. Assuming, as usually, that mutations arise as a Poisson process with rate $2N(\mu_R + \mu_P)$, then the rate of substitutions is

$$\begin{aligned}
\mathbb{E}(2N(\mu_R + \mu_P)\pi(s)) &= 2N(\mu_R + \mu_P)\mathbb{E}(\pi(s)) \\
&= 2N(\mu_R + \mu_P)\frac{1}{2N} \\
&= \mu_R + \mu_P
\end{aligned}$$

where we use earlier the fact that we computed the average fixation probability to be $\frac{1}{2N}$. So, since $(\mu_R + \mu_P)dt$ mutations will fix in the short time $dt$, we have that in in the short time $dt$, the change in mRNA is

$$\begin{aligned}
dR &= (\mu_R + \mu_P)dt\mathbb{E}(M_R \mid \text{fix}) \\
&= -2N\mu_R\sigma_R^2\left((1 - a_R c)\frac{R - a_R P - \theta_R}{V_R} + (c - a_p)\frac{P - a_P R - \theta_P}{V_P}\right) dt
\end{aligned}$$

where we've also approximated $2N - 1 \approx 2N$ for large N. Similarly, the change in protein is

$$\begin{aligned} dP &= (\mu_R + \mu_P) dt \mathbb{E}(M_P \mid \text{fix}) \\ &= -2N \left( \mu_R c \sigma_R^2 \left( (1 - a_R c) \frac{R - a_R P - \theta_R}{V_R} + (c - a_p) \frac{P - a_P R - \theta_P}{V_P} \right) \right. \\ &\quad \left. + \mu_P \sigma_P^2 \left( \frac{P - a_P R - \theta_P}{V_P} - a_R \frac{R - a_R P - \theta_R}{V_R} \right) \right) dt. \end{aligned}$$

Collecting terms and writing them as matrices, we get

$$F = \begin{bmatrix} 2N\mu_R\sigma_R^2 \left( \frac{1 - a_R c}{V_R} + \frac{a_P(a_P - c)}{V_P} \right) & -2N\mu_R\sigma_R^2 \left( \frac{a_R(1 - a_R c)}{V_R} + \frac{a_P - c}{V_P} \right) \\ c2N\mu_R\sigma_R^2 \left( \frac{1 - a_R c}{V_R} + \frac{a_P(a_P - c)}{V_P} \right) - 2N\mu_P\sigma_P^2 \left( \frac{a_R}{V_R} + \frac{a_P}{V_P} \right) & 2N\mu_P\sigma_P^2 \left( \frac{1}{V_P} + \frac{a_R^2}{V_R} \right) - c2N\mu_R\sigma_R^2 \left( -\frac{a_P - c}{V_P} - \frac{a_R(1 - a_R c)}{V_R} \right) \end{bmatrix}$$

and

$$\hat{\mathbf{X}} = \begin{bmatrix} \frac{\theta_R + a_R\theta_P}{1 - a_R a_P} \\ \frac{\theta_P + a_P\theta_R}{1 - a_R a_P} \end{bmatrix}.$$

Now, define

$$\begin{aligned} \alpha &= 2N\mu_R \frac{\sigma_R^2}{V_R} \\ \beta &= 2N\mu_R \frac{\sigma_P^2}{V_P} \end{aligned}$$

which are the rate of adaptation for mRNA and protein, respectively, and

$$\omega = \frac{V_R}{V_P}$$

which is the relative strength of selection on protein to mRNA ($\omega$ small means selection on mRNA is stronger, $\omega$ large means that selection on protein is stronger). Then, we can rewrite

$$\mathbf{F} = \begin{bmatrix} \alpha \left(1 - a_R c + a_P(a_P - c)\omega\right) & -\alpha \left(a_R(1 - a_R c) + (a_P - c)\omega\right) \\ \alpha c \left(1 - a_R c + a_P(a_P - c)\omega\right) - \beta \left(\frac{a_R}{\omega} + a_P\right) & \beta \left(1 + \frac{a_R^2}{\omega}\right) - \alpha c \left(a_R(1 - a_R c) + (a_P - c)\omega\right) \end{bmatrix}.$$

We also need to compute the matrix $\mathbf{\Sigma}$. Note that by our calculation of the (co)variance terms, and the rate of evolution argument, we have that

$$\mathbf{\Sigma}\mathbf{\Sigma}^T = \begin{bmatrix} \mu_R\sigma_R^2 & c\mu_R\sigma_R^2 \\ c\mu_R\sigma_R^2 & c^2\mu_R\sigma_R^2 + \mu_P\sigma_P^2 \end{bmatrix},$$

so that

$$\mathbf{\Sigma} = \begin{bmatrix} \sqrt{\mu_R}\sigma_R & 0 \\ c\sqrt{\mu_R}\sigma_R & \sqrt{\mu_P}\sigma_P \end{bmatrix},$$

For simplicity, define

$$\tau_R^2 = \mu_R \sigma_R^2$$

and

$$\tau_P^2 = \mu_P \sigma_P^2$$

so that

$$\mathbf{\Sigma} = \begin{bmatrix} \tau_R & 0 \\ c\tau_R & \tau_P \end{bmatrix},$$

which specifies our full model.

### 2 Constraining general model to test evolutionary hypotheses

We can take the previous model and consider two limits, to represent the extremes of mRNA dominated evolution and protein dominated evolution.

#### 2.1 mRNA-driven evolution

Here we assume that mutations that impact mRNA abundance have non-negligible impacts on protein abundance, and that mRNA abundance is the primary target of selection. Thus, we assume that $V_R \ll V_P$, and as such send $\omega \to 0$. We also set $a_R = 0$, to indicate that mRNA abundances do not experience selection related to protein abundances. Finally, we leave $a_P$ and $c$ to vary, to reflect how protein abundances are constrained by mRNA abundances and shaped by mRNA mutations, respectively. Applying these rules $\mathbf{F}$, $\hat{\mathbf{X}}$, and $\mathbf{\Sigma}$, we have

$$\mathbf{F} = \begin{bmatrix} \alpha & 0 \\ \alpha c - \beta a_P & \beta \end{bmatrix},$$

and

$$\hat{\mathbf{X}} = \begin{bmatrix} \theta_R \\ a_P \theta_R + \theta_P \end{bmatrix}$$

and

$$\mathbf{\Sigma} = \begin{bmatrix} \tau_R & 0 \\ c\tau_R & \tau_P \end{bmatrix}.$$

Under this model, the equilibrium covariance matrix, $\hat{\mathbf{V}}$, can be found by solving the system of equations

$$\hat{\mathbf{V}}\mathbf{F}^T + \mathbf{F}\hat{\mathbf{V}} = \mathbf{\Sigma}\mathbf{\Sigma}^T$$

to get

$$\hat{\mathbf{V}} = \begin{bmatrix} \frac{\tau_R^2}{2\alpha} & \left(c\frac{\alpha}{\alpha+\beta} + a_P \frac{\beta}{\alpha+\beta}\right)\frac{\tau_R^2}{2\alpha_R} \\ \left(c\frac{\alpha}{\alpha+\beta} + a_P \frac{\beta}{\alpha+\beta}\right)\frac{\tau_R^2}{2\alpha_R} & \frac{\tau_P^2}{2\beta} + \left(c^2\frac{\alpha}{\alpha+\beta} + a_P^2 \frac{\beta}{\alpha+\beta}\right)\frac{\tau_R^2}{2\alpha_R} \end{bmatrix}.$$

#### 2.2 Protein-driven evolution

In this model, we imagine that selection primarily acts on protein abundances, and selection on mRNA abundances primarily serves to keep it at the right level for a required protein abundance. Moreover, we assume that mutations that impact mRNA abundances have a negligible impact on protein abundances. To achieve a negligible impact of mRNA mutations on protein abundance, we first set $c = 0$. We also assume $V_P \ll V_R$ so that $\omega \to \infty$. Finally, we set $a_P = 0$ so that protein evolves toward its own optimum. Then,

$$\mathbf{F} = \begin{bmatrix} \alpha & -a_R\alpha \\ 0 & \beta. \end{bmatrix}$$

and

$$\hat{\mathbf{X}} = \begin{bmatrix} a_R\theta_P + \theta_R \\ \theta_P \end{bmatrix}$$

and

$$\mathbf{\Sigma} = \begin{bmatrix} \tau_R & 0 \\ 0 & \tau_P \end{bmatrix}.$$

As before, we can solve for the equilibrium covariance to get

$$\hat{\mathbf{V}} = \begin{bmatrix} \frac{\tau_R^2}{2\alpha} + a_R^2\frac{\alpha}{\alpha+\beta}\frac{\tau_P^2}{2\beta} & a_R\frac{\alpha}{\alpha+\beta}\frac{\tau_P^2}{2\beta} \\ a_R\frac{\alpha}{\alpha+\beta}\frac{\tau_P^2}{2\beta} & \frac{\tau_P^2}{2\beta} \end{bmatrix}.$$

#### 2.3 Independent evolution

In this model, protein and mRNA abundances evolve independently. Hence, we set all of the parameters that link them together to 0, namely $c = 0$, $a_R = 0$, and $a_P = 0$. Then,

$$\mathbf{F} = \begin{bmatrix} \alpha & 0 \\ 0 & \beta. \end{bmatrix}$$

and

$$\hat{\mathbf{X}} = \begin{bmatrix} \theta_R \\ \theta_P \end{bmatrix}$$

and

$$\mathbf{\Sigma} = \begin{bmatrix} \tau_R & 0 \\ 0 & \tau_P \end{bmatrix}.$$

Here, the equilibrium covariance is

$$\hat{\mathbf{V}} = \begin{bmatrix} \frac{\tau_R^2}{2\alpha} & 0 \\ 0 & \frac{\tau_P^2}{2\beta} \end{bmatrix}.$$

### 3 Validating phylogenetic model with simulations

To validate our implementation is working as expected, we generated numerous simulated datasets to test (1) the ability to identify the "correct" model using Deviance Information Criterion (DIC) and (2) to show that our model implementations can accurately estimate parameters averaged over 100s of "genes."

#### 3.1 Validating use of DIC to distinguish models

For these simulations, we generated 100 datasets, each with 100 genes. In this case, we assumed all genes evolved under the same sets of parameters i.e., there was no variation in the relevant parameters across genes. Data were simulated under both the protein-driven and mRNA-driven models with parameters being drawn from the following distributions:

$$\begin{aligned}
\alpha &\sim LogNorm(\mu_\alpha = \log\left(\frac{\log(2)}{(\text{Height}(T)/4)}\right), 1.5) \\
\beta &\sim LogNorm(\mu_\beta = \log\left(\frac{\log(2)}{(\text{Height}(T)/4)}\right), 1.5) \\
a_R, a_P &\sim Norm(\mu_{a_R, a_P} = 0.75, 0.25) \\
c &\sim Norm(\mu_c = 0.5, 0.5) \\
\tau_R &\sim LogNorm(\mu_{\tau_R} = -2.5, 1.5) \\
\tau_P &\sim LogNorm(\mu_{\tau_P} = -2.5, 1.5)
\end{aligned}$$

Here, Height$(T)$ represents the root-to-tip distance of the phylogenetic tree. Values of $\theta_R$ and $\theta_P$ for each gene $i$ were drawn from a normal distribution with mean 0 and standard deviation 1 to reflect that our gene expression values were converted to Z-scores for each species:

$$\theta_{R,i}, \theta_{P,i} \sim Norm(0, 1)$$

Due to the large number of MCMC chains being run, each chain was run for only 1,000 burn-in iterations with 10,000 adaptive iterations and 20,000 random-walk Metropolis-Hastings iterations, keeping every $5^{th}$ iteration. We kept only the model fits to a given simulated dataset where both the protein-driven and the mRNA-driven reached stationarity (as determined by **LaplacesDemon**). Regardless of the true model used for simulating the data, we observe a clear skew towards positive ΔDIC values, consistent with the true model being generally favored (Figure 1). This indicates that even with only 100 genes (less than 10% of the size of our dataset), we have no issue distinguishing between the two models.

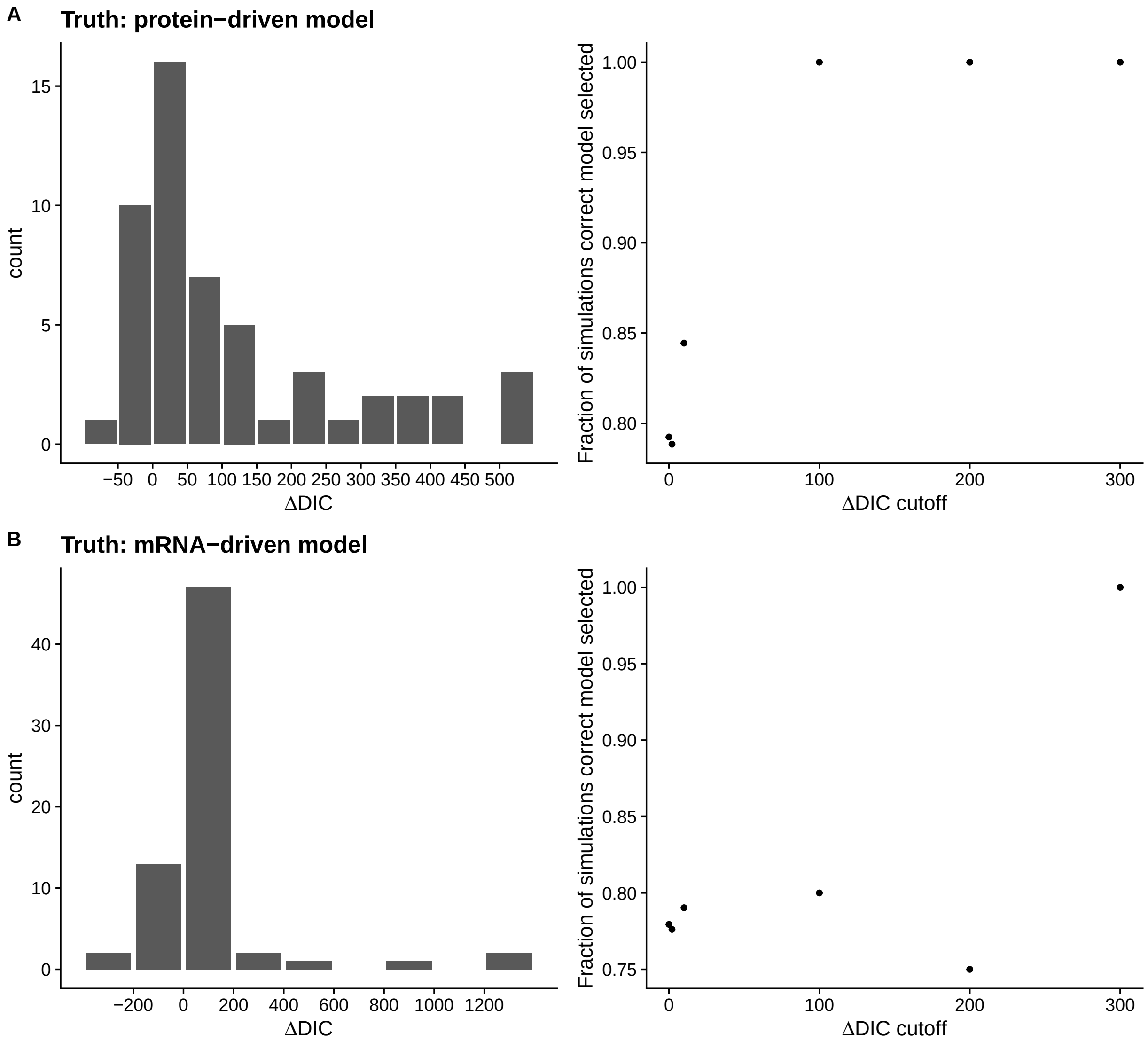

**Figure 1**: Assessing the ability of DIC to accurately distinguish the protein-driven and mRNA-driven model. $\Delta$DIC is calculated as $\text{DIC}_{\text{Wrong Model}}$ - $\text{DIC}_{\text{True Model}}$ such that $\Delta\text{DIC} > 0$ indicates the true model is the better of the two models. (A) Left panel shows the distribution of $\Delta$DIC values when the true model is the protein-driven model. Right panel shows the fraction of model comparisons in which the true model is correctly selected at a given $\Delta$DIC cutoff. (B) Panels are same as in (A), but when the mRNA-driven model is the true model.

#### 3.2 Validating ability of models to estimate population means of parameter

A key assumption of our model fitting strategy is that parameters $\alpha$, $\beta$, $a_R$, $a_P$, $\tau_R$, $\tau_P$, and $c$ are the same across all genes. This is unlikely to be the case. We reason that our parameter estimates reflect the population mean across our 1,641 genes. Instead of assuming that the parameters of interest were the same across all genes as done for model comparisons, we assumed parameters for the genes followed a distribution around a randomly chosen mean. As the mRNA-driven model is the most complex model under consideration here, we focused on our ability to estimate these parameters. Our MCMC implementation samples parameters $\alpha$, $\beta$, $\tau_R$, and $\tau_P$ are proposed on the log-scale and we sample population means to reflect this.

$$\begin{aligned}
\mu_\alpha, \mu_\beta &\sim Unif\left(\log\left(\frac{\log(2)}{(\text{Height}(T)/4)}\right), \log\left(\frac{\log(2)}{(2 \times \min(T))}\right)\right) \\
\mu_{\tau_R}, \mu_{\tau_P} &\sim Unif(-3, -2) \\
\mu_{a_P} &\sim Unif(0, 1) \\
\mu_c &\sim Unif(-1, 1)
\end{aligned}$$

As before, Height$(T)$ represents the root-to-tip distance of the phylogenetic tree and min$(T)$ represents the minimum branch length of all branches in the phylogenetic tree $T$. For each of the 100 simulations, each of which has a unique combination of population means for the parameters under consideration, we then drew the parameters for each of the 100 "genes" in our datasets according to the following distributions

$$\begin{aligned}
\alpha_i &\sim LogNorm(\mu_\alpha, 0.5) \\
\beta_i &\sim LogNorm(\mu_\beta, 0.5) \\
a_{P,i} &\sim Norm(\mu_{a_P}, 0.25) \\
c_i &\sim Norm(\mu_c, 0.25) \\
\tau_{R,i} &\sim LogNorm(\mu_{\tau_R}, 0.5) \\
\tau_{P,i} &\sim LogNorm(\mu_{\tau_P}, 0.5)
\end{aligned}$$

Values of $\theta_R$ and $\theta_P$ for each gene $i$ were drawn from a normal distribution with mean 0 and standard deviation 1 to reflect that our gene expression values were converted to Z-scores for each species:

$$\theta_{R,i}, \theta_{P,i} \sim Norm(0, 1)$$

As can be seen in Figure 2, parameter estimates from our model fits are generally in good agreement with the population mean used for simulating the data.

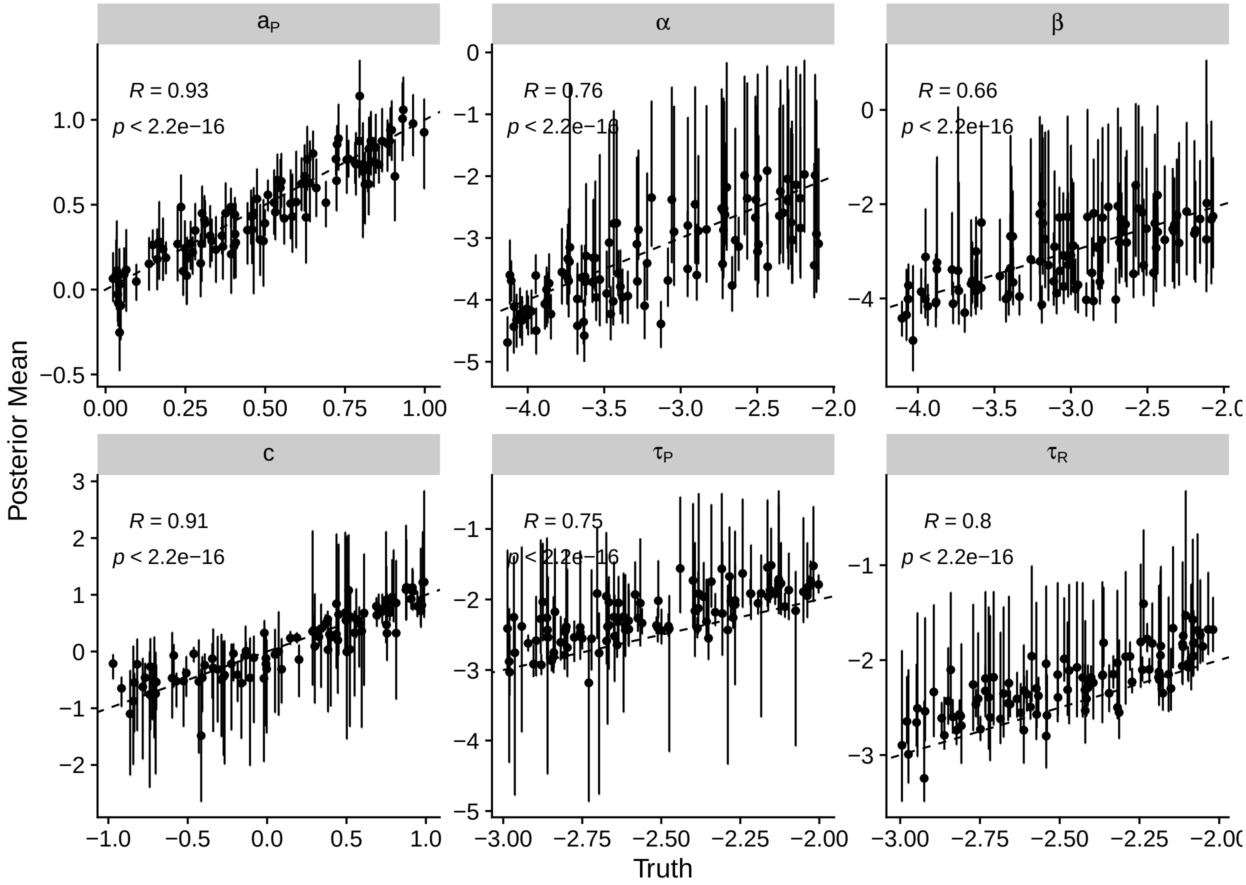

**Figure 2**: Comparing population means (on log scale for $\alpha$, $\beta$, $\tau_R$, and $\tau_P$ of simulated parameters to posterior estimates from mRNA-driven model fits (the true model). Dashed line represents the $y = x$ line. Error bars represent 95% credibility intervals. Spearman rank correlation $R$ and associated p-value are reported.

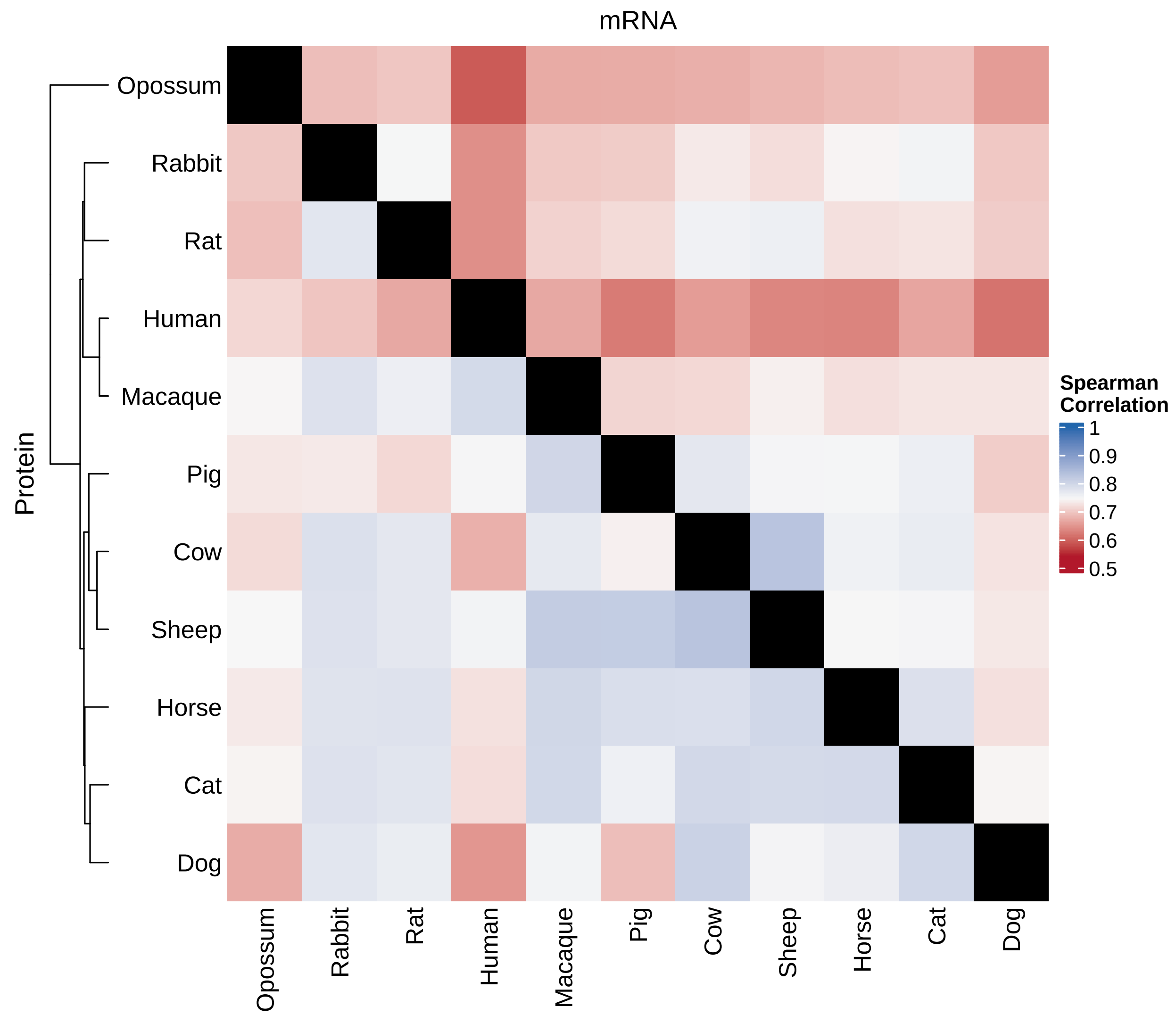

**Figure 3**: Heatmap showing across-species pairwise comparisons (measured as the Spearman rank correlation) of mRNA (upper triangle) and protein abundances (lower triangle). The phylogenetic tree of the 10 species is included on the lower triangle.

### 4 Empirical analyses

We obtained processed mRNA and protein abundances from [3]. The authors of [3] kindly provided the phylogenetic tree used for their analyses. We restricted our data set to the 1,641 one-to-one orthologous genes for which mRNA and protein abundances were available. mRNA and protein abundances were log-transformed and converted to Z-scores within each species for each replicate. The mean and standard error of the normalized expression values for each gene within each species were used as input into our model. We note that the human measurements from [3] had a relatively weak correlation with the other species (Figure 3). We suspect this is because the human samples were obtained from ATCC cell lines, while other samples were collected fresh [3].

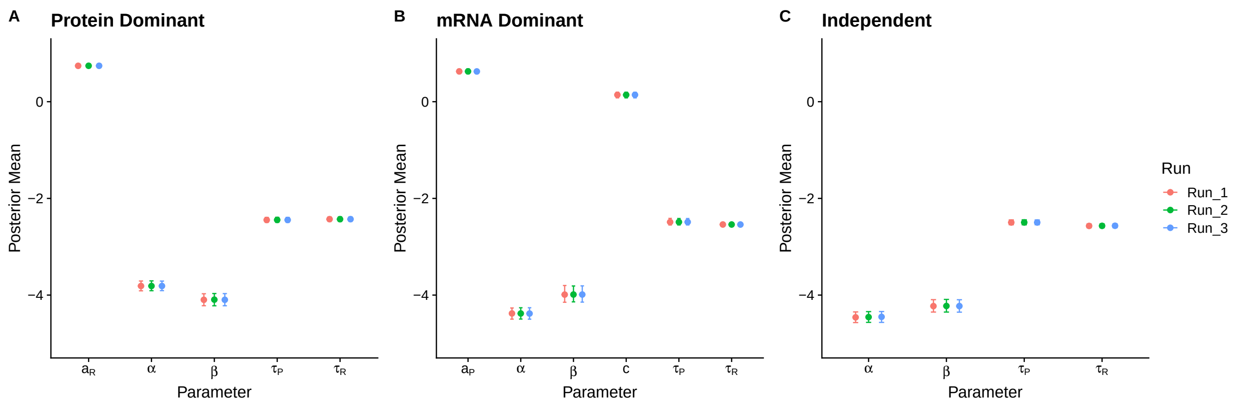

**Figure 4**: Comparing posterior means for parameters of interest across runs starting from different points in parameter space when applied to our real dataset. Error bars represent the 95% credible intervals.

#### 4.1 Fitting and comparing models

Models were fit using a Markov Chain Monte Carlo (MCMC) using code freom the R package **LaplacesDemon**. Local modifications of code from **LaplacesDemon** included code that sped up calculations of our likelihoods, as well as minor bug fixes. Parameters were proposed and accepted using a blockwise scheme. We first ran with a burn-in of 1000 iterations, followed by a robust adaptive metropolis adaptive MCMC [4] to tune parameter fitting using an acceptance target ratio of 0.44. After the adaptive MCMC, a standard random walk Metropolis-Hastings chain was run for parameter estimation. We found that multiple runs of the MCMC starting at different points in parameter space converged for all three models (protein-driven, mRNA-driven, and independent) presented here. The MCMCs were run until they (1) achieved stationarity and (2) had at least 100 effective samples for each parameter of interest. Likelihoods were calculated using **PCMBase** [5]. We note 3 independent runs of our MCMCs, each starting from different points in parameter space, converged on the same parameter estimates (Figure 4).

#### 4.2 Testing predictions with functional genomic data

##### 4.2.1 Stratifying by gene essentiality

Genes that are "essential" to an organism (i.e., a functional form of the gene is necessary for viability) are thought to be subject to greater evolutionary constraint. Here, we tested this hypothesis in the context of gene expression using genes previously identified as essential [6]. These classifications were based on 1,085 CRISPR knockout screens quantifying the effects of genes on cell proliferation and survival obtained from [6]. For our study, genes were classified as "essential" as in [6]; all other genes were classified as "non-essential." The protein-driven model was fit to each set of genes independently to test if essential genes exhibit unique patterns in gene expression evolution.

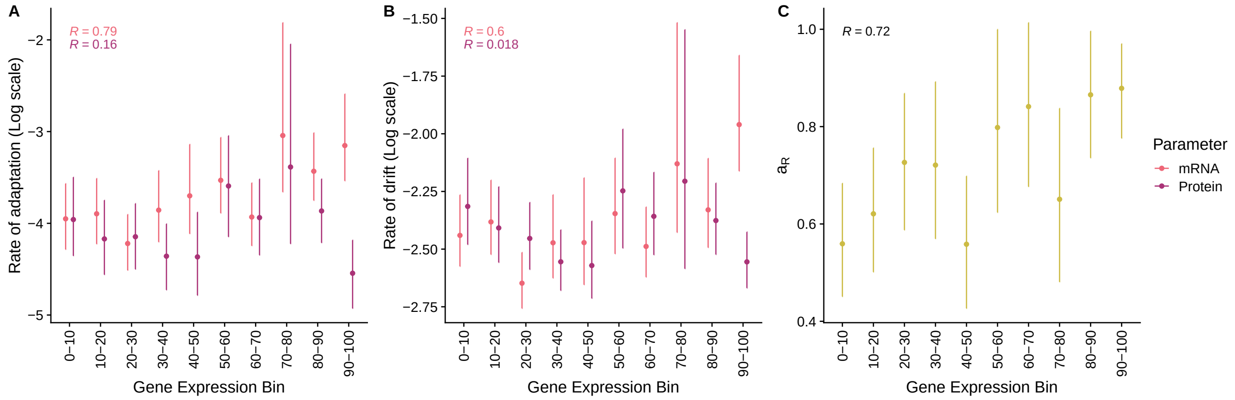

**Figure 5**: Variation in parameters across gene expression bins based on human protein abundances data taken from PaxDb [7].

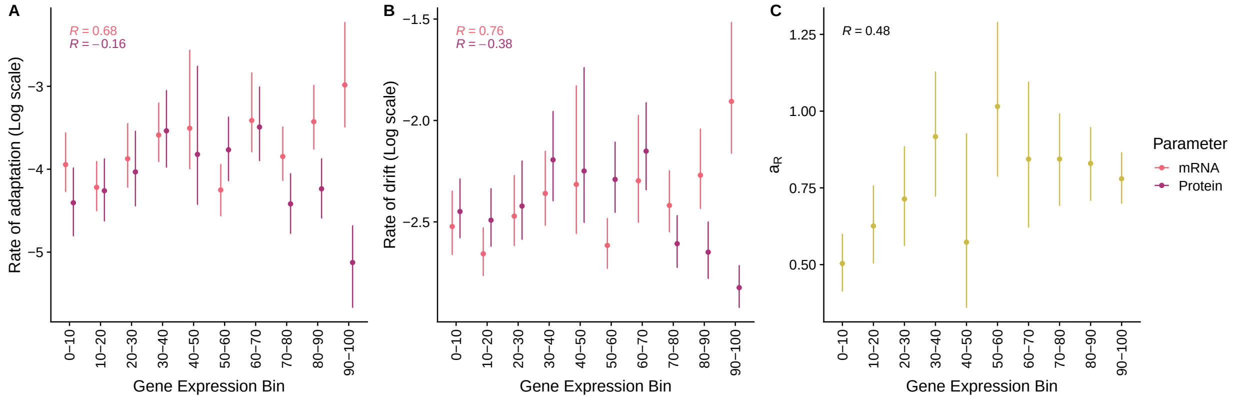

**Figure 6**: Variation in parameters across gene expression bins based on mean protein abundances from the species used in this study.

###### 4.2.2 Stratifying by gene expression abundances

To test for the effects of gene expression on the co-evolutionary dynamics between mRNA and protein abundances, we divided genes into bins representing the deciles of protein abundances. To avoid fitting our models to data used to generate the bins, we obtained independent protein abundances from PaxDB [7] for humans, an out-of-sample species. Specifically, we used the integrated whole-organism dataset. We pulled the 1,641 orthologous genes under consideration from the independent human data and binned them into deciles based on the integrated protein abundances, creating 9 bins with 164 genes and 1 with 165 genes. Our simulations suggested 100 genes were sufficient to capture the population means of the parameters of interest. We re-fit the protein-driven model to the mRNA and protein abundances for each bin, allowing us to examine how the rate of adaptation and drift vary as a function of gene expression (Figure 5). We performed a similar analysis using the mean protein abundances from the species under consideration, finding consistent results (Figure 6).

Many further analyses are based on the gene expression decile bins from the independent human protein abundance data. In Figure 7 and Figure 8, we show the relationship between gene expression and these features, finding they are generally weak.

##### 4.2.3 Determining mutational input using the number of regulatory elements and eQTLs

The underlying regulatory structure of a gene is a key aspect shaping the evolution of its expression levels. Regulatory complexity, particularly the number of enhancers, correlates with macroevolutionary patterns of mRNA abundance evolution [8]. A greater number of regulatory elements may confer some degree of mutational robustness, but also increase the probability that a mutation (whether it be harmful or beneficial) occurs in a regulatory element [9]. Predicted enhancers were identified in human keratinocyte cells – a cell-type found in the epidermis – using the EnhancerAtlas database [10, 11]. The number of enhancers was calculated for each gene. Using our decile bins from the gene expression analysis, we then calculated the mean number of enhancers per gene per bin. A similar analysis was repeated using enhancers in mouse kidney tissue (same lineage, different tissue type) and *D. melanogaster* larvel cell lines (Figure 9).

As an alternative potential proxy of mutational input, we used the effect sizes of human expression quantitative trait loci (eQTLs) taken from the Gene Tissue-Expression (GTEx) database [12, 13]. As we are not interested in the direction of the effect, we calculated the mean absolute effect size for each gene across all available samples in the GTEx. Similar to the enhancer analysis, we then calculated the mean absolute effects size per gene per bin based on our *a priori* determined gene expression decile bins based on the independent human protein abundances.

##### 4.2.4 Quantifying gene properties related to mRNA translation and translation efficiency

Changes to mRNA translation are thought to play a role in shaping protein abundances independent of mRNA abundances, although translation can impact mRNA degradation rates [14]. We tested if our parameters generally reflected the evolution of translation using four different approaches. Two of these approaches constitute the evolution of gene properties thought to reflect the evolution of protein abundances via changes to mRNA translation, while the other two concern the evolution of translation efficiency itself.

*Gene properties related to protein abundance adaptation*

First, we tested if divergences between human and mouse mRNA secondary structure (specifically, the ensemble free energy) near the protein-coding sequence start codon were correlated with the rate of adaptation of protein abundances. Previous work concluded changes to mRNA secondary structure around the site of translation initiation can significantly impact protein abundances [15]. Computational estimates of mRNA secondary structure in human and mouse were taken from [16], who estimated the ensemble free energies of 35 nucleotide windows in the region 50 nucleotides upstream and downstream of the start codon, resulting in 101 estimates of mRNA secondary structure around the start codon. Absolute divergences were calculated from the free energies for each orthologous gene in each 35-nucleotide window between human and mouse. For each of our decile gene expression bins, the median divergence was calculated and compared to the rates of adaptation for both mRNA ($\alpha$) and protein ($\beta$) abundances.

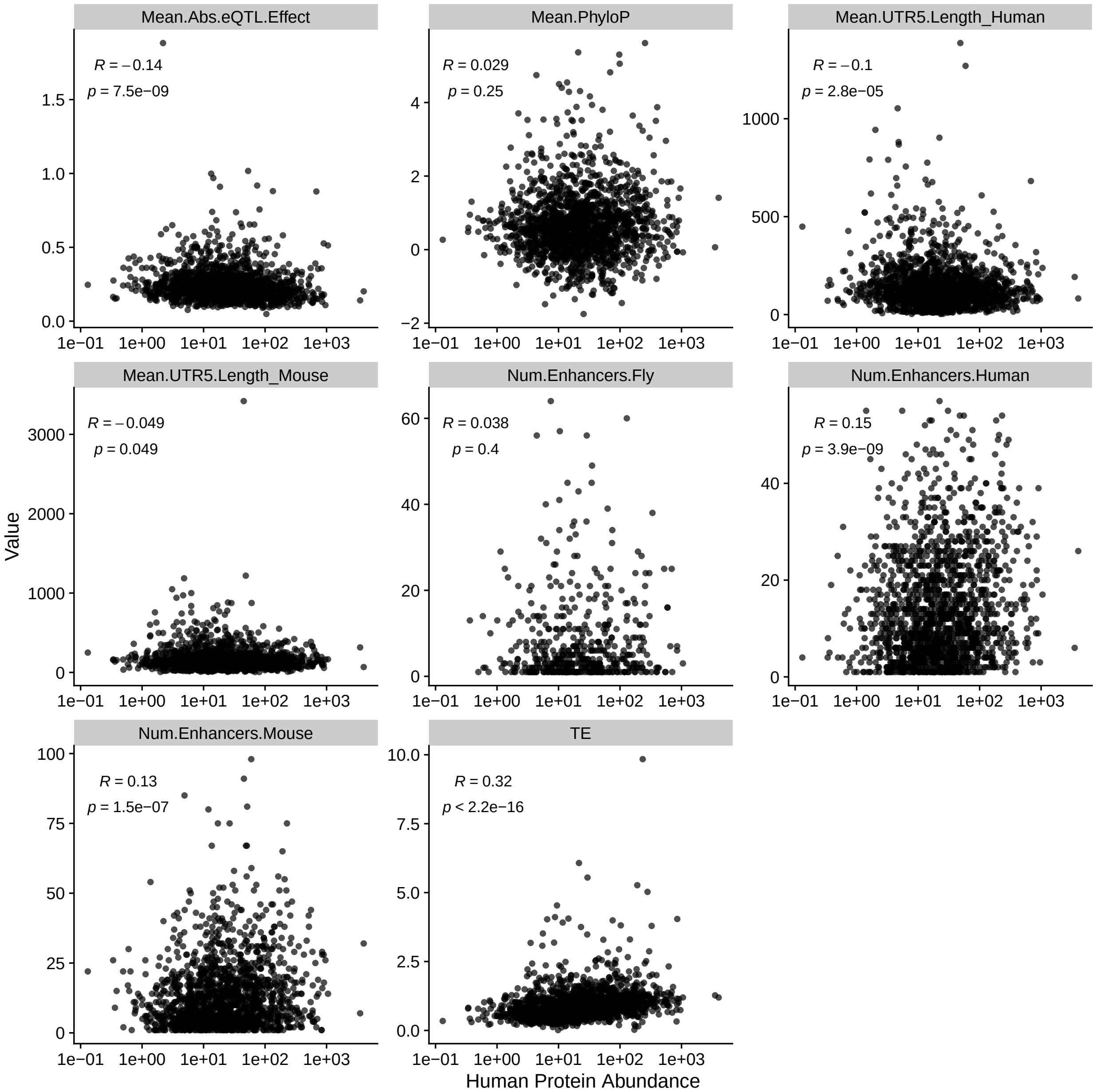

**Figure 7**: Relationship between human independent protein abundances from PaxDB [7] and numerous features related to gene expression evolution. TE indicates translation efficiency. Spearman rank correlations $R$ are reported.

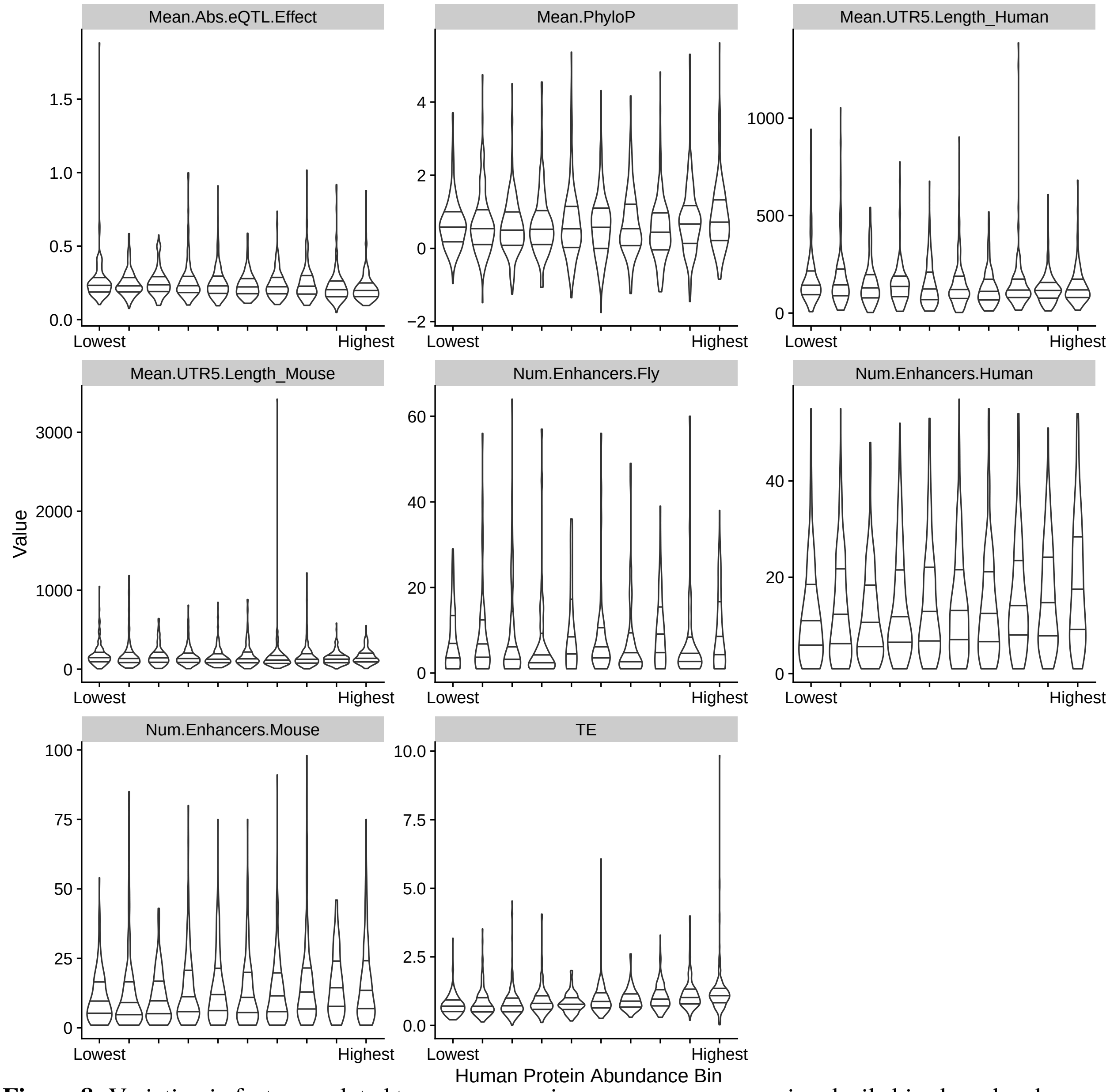

**Figure 8**: Variation in features related to gene expression across gene expression decile bins based on human independent protein abundances from PaxDB [7]. TE indicates translation efficiency. Lines going across the violin plots indicate the interquartiles.

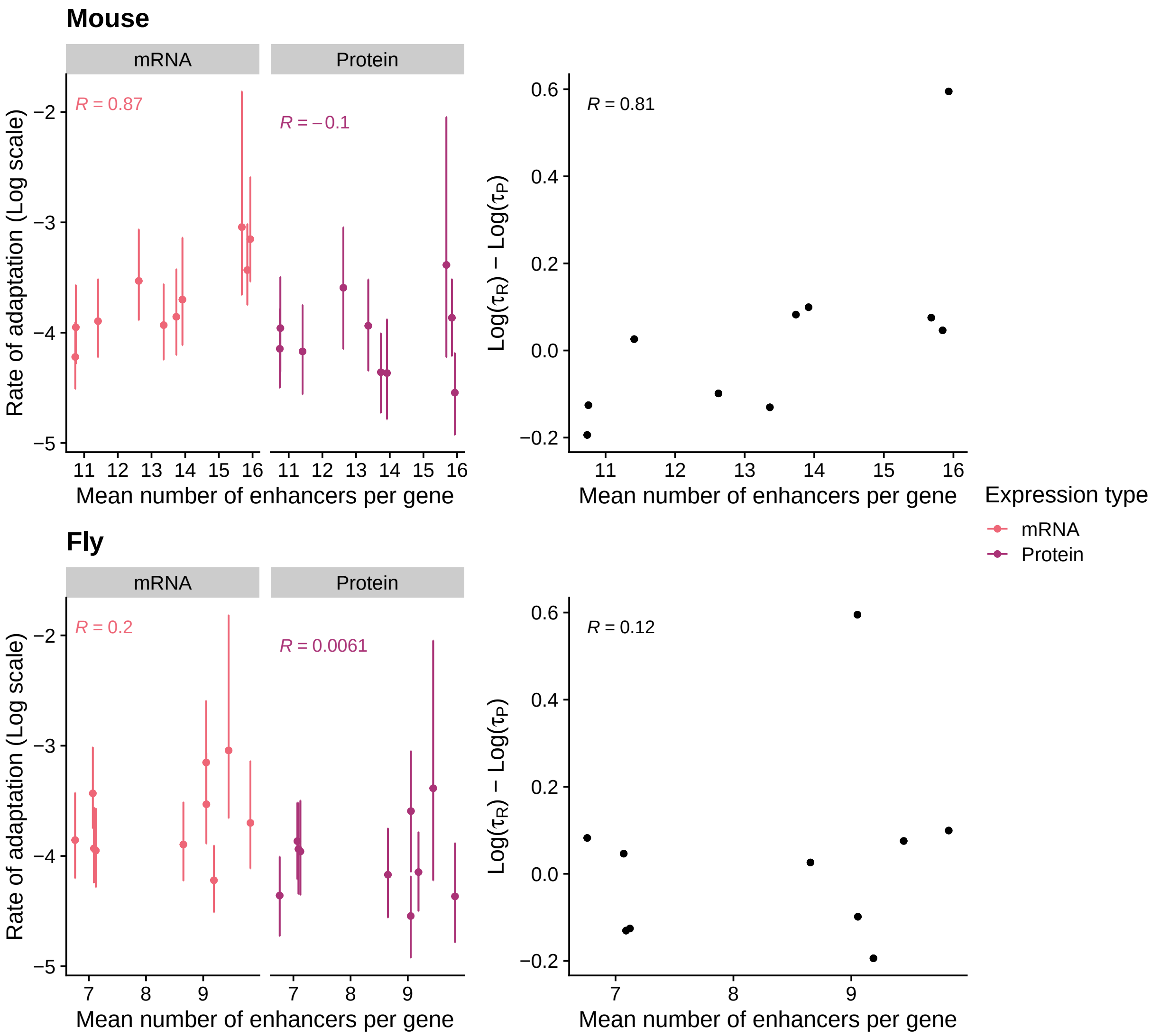

**Figure 9**: Comparison of rate of adaptation parameters $\alpha$ (mRNA) and $\beta$ (protein) and rate of drift parameters $\tau_R, \tau_P$ to mean number of enhancers in from mouse kidney (top) and fly larval cells.

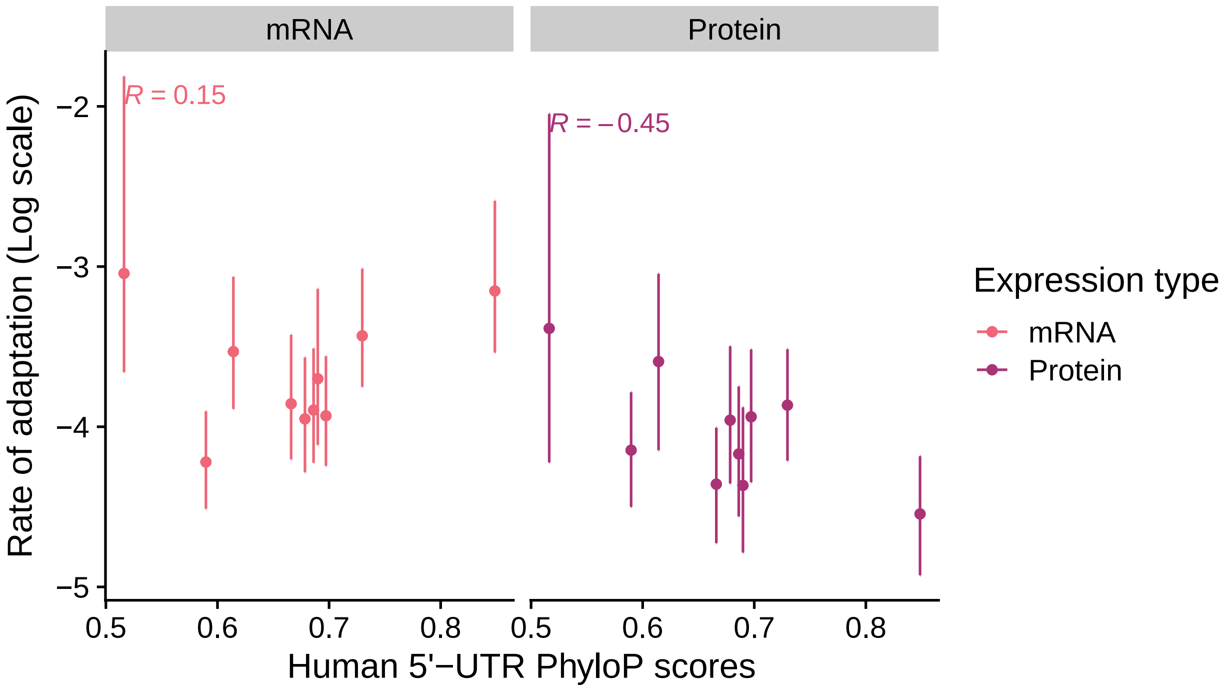

**Figure 10**: Comparison of rate of adaptation parameters $\alpha$ (mRNA) and $\beta$ (protein) to per-bin average PhyloP scores estimated for 5'-UTR regions taken from [17]. Spearman rank correlation $R$ is reported. Bins were based on independent human protein abundance data taken from PaxDB [7].

Second, we tested if the overall sequence conservation of the 5' untranslated region (5'-UTR) was correlated with the rate of adaptation of protein abundances. As changes to the 5'-UTR region – a key *cis* regulator of mRNA translation – can impact translation, genes showing higher rates of adaptation to protein abundances are expected to show less conservation of the 5'-UTR. We obtained PhyloP scores from [17], which calculated per-gene average PhyloP scores for human 5'-UTRs relative to vertebrates. Briefly, positive PhyloP scores indicate a nucleotide or region is more conserved than expected under neutrality (i.e., negative or purifying selection), while negative PhyloP scores indicate a nucleotide or region that is more diverged than expected under neutrality (i.e., positive or diversifying selection). Calculating the average PhyloP scores per bin, we found that the rate of adaption of protein abundances was negatively correlated with the human 5'-UTR PhyloP scores. In contrast, the 5'-UTRs PhyloP scores are positively correlated with the rates of adaptation for mRNA abundances. Taken together, these results suggest genes exhibiting less conservation of 5'-UTRs adapting faster at the level of protein abundances with little change to mRNA abundances.

*Evolution of translation efficiency*

In our best model, the optimal mRNA and protein abundances are linked by the parameter $a_R$, such that $a_R$ may reflect factors related to translation efficiency. We tested if our estimates of the optimal relationship between our mRNA and protein abundances as mediated by $a_R$ agree with common empirical estimate of translation efficiency from Ribo-seq (a.k.a. ribosome profiling) and RNA-seq data, specifically the per-gene ratio of mapped Ribo-seq reads to RNA-seq reads. We obtained human Ribo-seq and RNA-seq transcripts per million (TPM) estimates from [18], from which we calculated $\text{Log}(\text{TE}_g) = \text{Log}\left(\frac{\text{Ribo TPM}}{\text{RNA TPM}}\right)$ for each

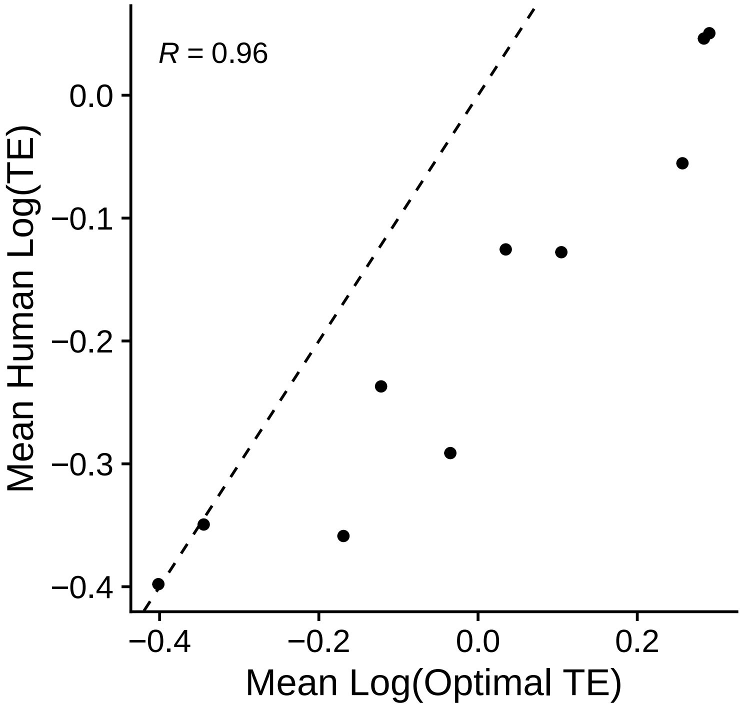

**Figure 11**: Comparison of optimal translation efficiency (as estimated via the protein selection strong model) and empirical estimates of translation efficiency for the human transcriptome [18]. Bins were based on independent human protein abundance data taken from PaxDB [7]. The Spearman rank correlation $R$ is reported. The dashed line indicates $y = x$.

gene $g$. To test if our parameters generally reflect these translation efficiencies, we compared the empirical translation efficiencies to the "optimal" translation efficiencies estimated by our model. We note that our estimates of per-gene optima $\theta_R + a_R\theta_P$ and $\theta_P$ are on the log-scale, such that we calculate the optimal translation efficiency (OTE) for each gene as $\text{Log(OTE}_g) = \theta_{P,g} - (\theta_{R,g} + a_R\theta_{P,g})$ for each gene $g$. To stay consistent with our other analyses, we tested if the per-bin average empirical TEs were correlated with our per-bin average optimal TEs. We observed a clear correlation ($R = 0.96$) between the empirical and optimal TEs, indicating our model generally captures the relationship between protein and mRNA abundances as modulated by mRNA translation. This suggests $a_R$ largely reflects factors related to translation efficiency.

As stated above, the 5'-UTR is an important *cis* regulator of mRNA translation. Thus, we might expect $a_R$ to be correlated with properties of the 5'-UTR. 5'-UTR widths for genes annotated as level = 1 in the GFF3 file were obtained for human and mouse transcripts downloaded from Gencode for genome version GRCh38.p14 and GRCm39 (reference chromosome), respectively. For genes with multiple isoforms, the mean length of the 5'-UTR was used. The mean 5'-UTR length was calculated per gene expression bin and correlated with the $a_R$ parameters for each bin. Consistent with $a_R$ reflecting factors related to translation efficiency, we see a clear negative correlation between $a_R$ and the 5'-UTR length in both human and mouse (Spearman rank correlation $R = -0.62$ and $R = -0.64$, respectively).

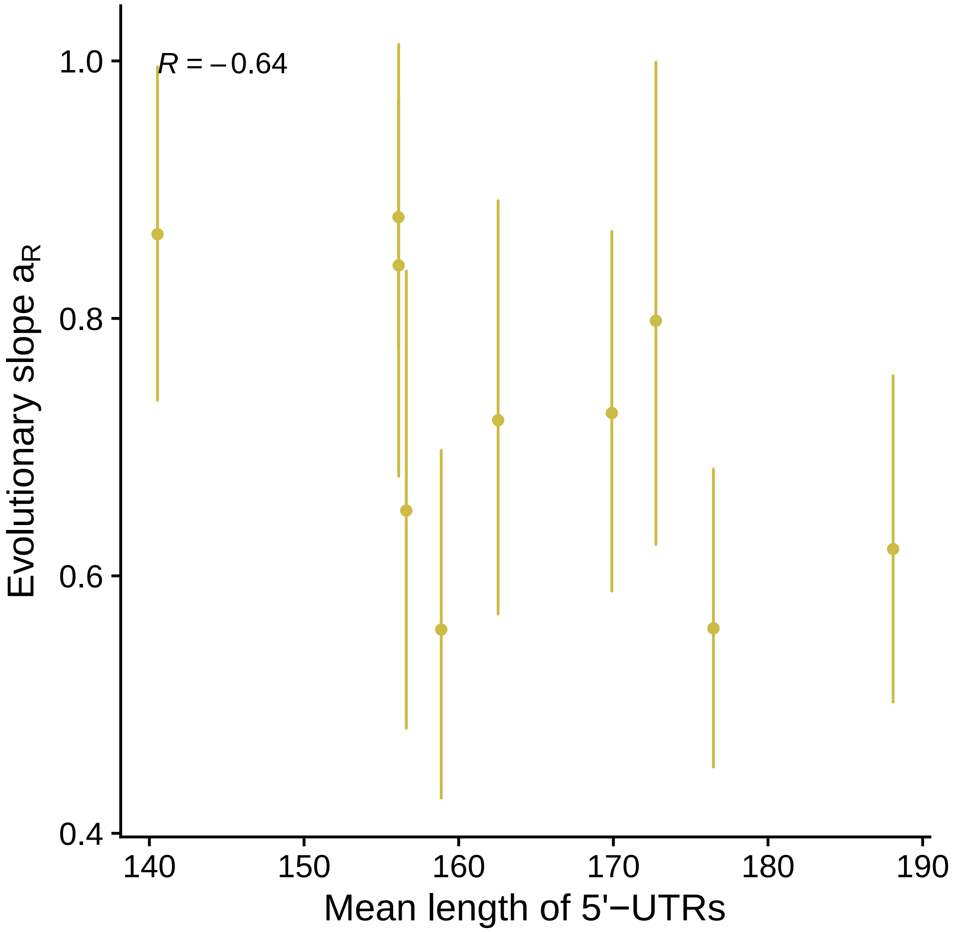

**Figure 12**: Comparison of $a_R$ and the per-bin mean 5'-UTR length in mouse. Bins were based on independent human protein abundance data taken from PaxDB [7]. The Spearman rank correlation $R$ is reported

[12] Aguet, F, Barbeira, A. N, Bonazzola, R, Brown, A, Castel, S. E, Jo, B, Kasela, S, Kim-Hellmuth, S, Liang, Y, Oliva, M, Flynn, E. D, Parsana, P, Fresard, L, Gamazon, E. R, Hamel, A. R, He, Y, Hormozdiari, F, Mohammadi, P, Muñoz-Aguirre, M, Park, Y. S, Saha, A, Segrè, A. V, Strober, B. J, Wen, X, Wucher, V, Ardlie, K. G, Battle, A, Brown, C. D, Cox, N, Das, S, Dermitzakis, E. T, Engelhardt, B. E, Garrido-Martín, D, Gay, N. R, Getz, G. A, Guigó, R, Handsaker, R. E, Hoffman, P. J, Im, H. K, Kashin, S, Kwong, A, Lappalainen, T, Li, X, MacArthur, D. G, Montgomery, S. B, Rouhana, J. M, Stephens, M, Stranger, B. E, Todres, E, Viñuela, A, Wang, G, Zou, Y, Anand, S, Gabriel, S, Graubert, A, Hadley, K, Huang, K. H, Meier, S. R, Nedzel, J. L, Nguyen, D. T, Balliu, B, Conrad, D. F, Cotter, D. J, deGoede, O. M, Einson, J, Eskin, E, Eulalio, T. Y, Ferraro, N. M, Gloudemans, M. J, Hou, L, Kellis, M, Li, X, Mangul, S, Nachun, D. C, Nobel, A. B, Park, Y, Rao, A. S, Reverter, F, Sabatti, C, Skol, A. D, Teran, N. A, Wright, F, Ferreira, P. G, Li, G, Melé, M, Yeger-Lotem, E, Barcus, M. E, Bradbury, D, Krubit, T, McLean, J. A, Qi, L, Robinson, K, Roche, N. V, Smith, A. M, Sobin, L, Tabor, D. E, Undale, A, Bridge, J, Brigham, L. E, Foster, B. A, Gillard, B. M, Hasz, R, Hunter, M, Johns, C, Johnson, M, Karasik, E, Kopen, G, Leinweber, W. F, McDonald, A, Moser, M. T, Myer, K, Ramsey, K. D, Roe, B, Shad, S, Thomas, J. A, Walters, G, Washington, M, Wheeler, J, Jewell, S. D, Rohrer, D. C, Valley, D. R, Davis, D. A, Mash, D. C, Branton, P. A, Sobin, L, Barker, L. K, Gardiner, H. M,

Mosavel, M, Siminoff, L. A, Flicek, P, Haeussler, M, Juettemann, T, Kent, W. J, Lee, C. M, Powell, C. C, Rosenbloom, K. R, Ruffier, M, Sheppard, D, Taylor, K, Trevanion, S. J, Zerbino, D. R, Abell, N. S, Akey, J, Chen, L, Demanelis, K, Doherty, J. A, Feinberg, A. P, Hansen, K. D, Hickey, P. F, Hou, L, Jasmine, F, Jiang, L, Kaul, R, Kibriya, M. G, Li, J. B, Li, Q, Lin, S, Linder, S. E, Pierce, B. L, Rizzardi, L. F, Smith, K. S, Snyder, M, Stamatoyannopoulos, J, Tang, H, Wang, M, Branton, P. A, Carithers, L. J, Guan, P, Koester, S. E, Little, A. R, Moore, H. M, Nierras, C. R, Rao, A. K, Vaught, J. B, & Volpi, S. (2020) *Science* **369**, 1318–1330.

[13] Carithers, L. J, Ardlie, K, Barcus, M, Branton, P. A, Britton, A, Buia, S. A, Compton, C. C, Deluca, D. S, Peter-Demchok, J, Gelfand, E. T, Guan, P, Korzeniewski, G. E, Lockhart, N. C, Rabiner, C. A, Rao, A. K, Robinson, K. L, Roche, N. V, Sawyer, S. J, Segrè, A. V, Shive, C. E, Smith, A. M, Sobin, L. H, Undale, A. H, Valentino, K. M, Vaught, J, Young, T. R, Moore, H. M, Barker, L, Basile, M, Battle, A, Boyer, J, Bradbury, D, Bridge, J. P, Brown, A, Burges, R, Choi, C, Colantuoni, D, Cox, N, Dermitzakis, E. T, Derr, L. K, Dinsmore, M. J, Erickson, K, Fleming, J, Flutre, T, Foster, B. A, Gamazon, E. R, Getz, G, Gillard, B. M, Guigó, R, Hambright, K. W, Hariharan, P, Hasz, R, Im, H. K, Jewell, S, Karasik, E, Kellis, M, Kheradpour, P, Koester, S, Koller, D, Konkashbaev, A, Lappalainen, T, Little, R, Liu, J, Lo, E, Lonsdale, J. T, Lu, C, MacArthur, D. G, Magazine, H, Maller, J. B, Marcus, Y, Mash, D. C, McCarthy, M. I, McLean, J, Mestichelli, B, Miklos, M, Monlong, J, Mosavel, M, Moser, M. T, Mostafavi, S, Nicolae, D. L, Pritchard, J, Qi, L, Ramsey, K, Rivas, M. A, Robles, B. E, Rohrer, D. C, Salvatore, M, Sammeth, M, Seleski, J, Shad, S, Siminoff, L. A, Stephens, M, Struewing, J, Sullivan, T, Sullivan, S, Syron, J, Tabor, D, Taherian, M, Tejada, J, Temple, G. F, Thomas, J. A, Thomson, A. W, Tidwell, D, Traino, H. M, Tu, Z, Valley, D. R, Volpi, S, Walters, G. D, Ward, L. D, Wen, X, Winckler, W, Wu, S, Zhu, J, Abdallah, A, Addington, A, Anderson, J. M, Bender, P. K, Cosentino, M, Diaz-Mayoral, N, Engel, T, Garci, F, Green, A, Hammond, T, Jaffe, K, Keen, J, Kennedy, M, Kigonya, P, Lander, B, Nampally, S, Ny, C, Robb, J, Santhanum, V, Sharopova, N, Singh, S, Soria, C, Sturcke, A, Sukari, S, Thomson, E. J, Tomaszewski, M, Trowbridge, C, Udoye, F, Vanscoy, D, Vatanian, N, Wilder, E. L, & Williams, P. (2015) *Biopreservation and Biobanking* **13**, 311–317.
